# Semi-automated annotation refinement accelerates cell type identification in brain spatial and single-cell studies

**DOI:** 10.64898/2026.07.30.741795

**Authors:** Dominic J. Acri, Richard Mustaklem, Liam Horan-Portelance, Luke C. Dabin, Jung Hyun Park, Kelly A. Hartigan, Holly N. Kersey, Monica E. Mesecar, J. Raphael Gibbs, Mark R. Cookson, Jungsu Kim

## Abstract

**Background:** single-cell and spatial omic techniques have enabled the investigation of cell type specific alterations in biologically complex tissues. In an effort to map cell taxonomies, large atlas-based studies and multi-laboratory consortia have created sets of annotated cell types. However, application of atlas- or database-level knowledge to individual studies is often resource-limited and computational demands scale with the size of both query and reference datasets.

**Results:** Here, we report a statistical framework for rapid label transfer using summary statistics and user-defined hyperparameters. Semi-Automated Hand Annotation (SAHA)^1^ allows the user to investigate magnitude, directionality, and statistical significance of matches between unnamed query clusters and reference cell types using either marker-based or marker-free methodologies. By pre-loading the package with summary statistics from the Allen Brain Cell Atlas of the mouse brain, the SAHA R package is capable of rapid cell type comparisons that closely mimic cell typing by integration-based annotation strategies. Furthermore, this flexible package is capable of comparisons across omic modalities, cluster resolutions, and annotations from any study where summary statistics are available. We demonstrate this flexibility by using multiple single-nuclei studies of the mouse cerebellum, mouse cerebral cortex, human cerebral cortex, human peripheral blood mononuclear cells, and one mouse spatial transcriptomic assay. Importantly, this method avoids privacy concerns as it does not require the sharing or deposition of raw data in a web-based tool.

**Conclusions:** As a result, SAHA offers a non-deterministic annotation reporting structure with automated html reports and summary statistics for transparency in cell typing decisions. Taken together, this scalable framework implemented as a package in R affords increased biological insight into the annotation of single-cell and spatial datasets.

**SHORT SUMMARY:** Acri and colleagues present rapid cell type annotation without the need for dataset integration. This paper outlines the utility of the package, SAHA, in annotating neurological datasets.

## INTRODUCTION

Technologies focused on assaying transcriptomes, proteomes, and epigenomes with single-cell precision or spatial resolution have increased the granularity and dimensionality with which cell types can be classified^2–4^. Historically, cell type identity has been defined through a combination of histological staining^5^, morphological analysis^6^, and marker profiling^7^. The addition of transcriptome-, proteome-, and genome-wide quantifications have reinforced previous identity descriptors while also enabling the discovery of substates otherwise masked during lower throughput assays. A major challenge that comes with this wealth of new information is the growing scale of datasets and the proportional increase in computational costs. This challenge is especially critical in complex biological tissues such as the central nervous system, where certain neuronal subtypes have been shown to be selectively vulnerable to neurodegenerative diseases^8^ and glial substates are implicated in critical immune responses to pathology^9^. As many computational pipelines are focused on deterministic annotations (i.e. “What is this cell type most likely to be?”), little work has been done to enable rapid cross-study comparisons (i.e. “How much does my cell type A look like another study’s cell type B?”).

Many features that drive identity have been shown to poorly generalize across omic modality and assay technique^10^. For example, microglia marker IBA1 is one of the most commonly used protein markers in histological staining^11^. However, the gene which encodes IBA1, AIF1, does not necessarily translate into a reliable transcriptomic marker of microglia in single-nuclei RNA sequencing (snRNA-seq) studies^12^. Whether this discrepancy is due to the known discordance between proteomic and transcriptomic assays^13^ or a true divergence between protein- and RNA-markers of cell identity is unknown. These challenges necessitate the creation of new assay-specific ground truth on cell type identity. Current solutions include the creation of consensus atlases^14–20^ and brute-force databases^21,22^ containing all possible cell type designations and markers-of-interest. Many broad^14,15^, organ-specific^19,20^, or disease specific databases^23^ allow researchers to access which markers are present for a given cell type. However, annotating new sample series based on select markers and without a statistical framework is prone to misannotation, over-clustering, inconsistent naming strategies, or inappropriately merged subclusters.

The practical consequence of this is that researchers are left making annotation decisions without adequate reporting. Even with the advent of integration-based label transfer^24^, cell cluster determination is still sensitive to study-specific artifacts and annotation disagreements. Technical artifacts such as ambient RNA, heterotypic doublets, and high mitochondrial read content have been shown to obscure cell type identity^25^. Other artifacts such as sample preparation or donor-specific variability pose similar clustering challenges^26,27^. Minimizing these artifacts is especially critical with recent trends in the discovery of cellular substates or fate trajectories^28^. To enable reproducible and statistical frameworks for cell annotation there is a need for rapid cross-study comparisons that can be iteratively run across multiple quality control or alternative clustering pipelines.

Because of this gap in available methodology for cell type annotation, we aimed to create an R package based on cluster marker genes and per-cluster average expression profiles. We created the Semi-Automated Hand Annotation (SAHA) R package which implements marker-based (cluster markers) and marker-free (average expression profiles) statistical analyses between a query dataset and any user-defined reference dataset. Importantly, since the pipeline only requires two dataframes–one from the query and one from the reference–this solution can scale up in terms of donor and cells without a concordant increase in compute resources. To demonstrate the utility of SAHA, we performed annotations of publicly available mouse single-cell and spatial datasets against the Allen Brain Cell (ABC) Atlas data preloaded within the package. We found that SAHA either matched or improved upon the existing level of annotation. Furthermore, we demonstrate the flexibility of SAHA by performing glial substate annotation from the human brain of Alzheimer’s disease patients^29^, and human peripheral blood mononuclear cells^30^. The application of user-defined reference databases resulted in reproducible, descriptive reports that enable biologically-driven annotation decisions. Taken together, SAHA provides a statistical framework to increase transparency and reproducibility in cell type annotation across the mammalian brain and beyond.

## RESULTS

The “Whole adult mouse brain taxonomy of cell types” (ABC Atlas) is a complete taxonomy of cell types in the mouse brain created through 10x Genomics scRNA-seq expression data^20^. At the level defined by the Allen Institute as “subclass” (e.g. 311 CBX MLI1 Megf11 Gaba, 314 CB Granule Glut, 317 Astro CB NN), we calculated marker genes and average expression profiles. These dataframes, along with the accompanying metadata from the original study, were accessed throughout all comparisons to mouse brain data through the SAHAdata package^31^. The final taxonomy available as part of SAHA included 338 unique cell types (subclass) organized into 34 broad cell type groupings (class) and 3 macro-level groupings (neurotransmitter) present across 12 regions of the mouse brain (**See Methods**). There was an average of 52.08 +/- 23.35 subclasses present (mean +/- sd, range: 19-98) per region.

### Query-database comparisons leverage similarity scoring for user-based annotation decisions

To investigate the efficacy of SAHA in identifying transcriptionally distinct major cell classes, we obtained a processed snRNA-seq dataset (611,034 nuclei, 6 mice ranging from ages E18-P60) from microdissected mouse cerebellum^32^. Using the cell annotations published in the original manuscript (e.g.. Granule, MLI1, Astrocyte), we calculated broad cell type marker genes, average expression profiles, and top 2,000 variable features used for dimensionality reduction (**Fig. 1A**, **Table S1-2**). Across 18 annotated clusters there were 637.11 +/- 314.92 (mean +/- sd, range: 232-1,416) markers per cluster and 24,409 genes with at least one non-zero count value. We then tested the performance of both marker-based and marker-free approaches. Using the SAHA marker-based pipeline, we compared markers from the Kozareva_2021 dataset to the ABC atlas subclasses within the cerebellum (CB, **Figs. 1B, Supp Data 1-2**). To perform the SAHA marker-free pipeline, the same comparison was made for expression profiles of the 2,000 variable features found in the Kozareva_2021 dataset (**Figs. 1C**).

**Figure 1.**
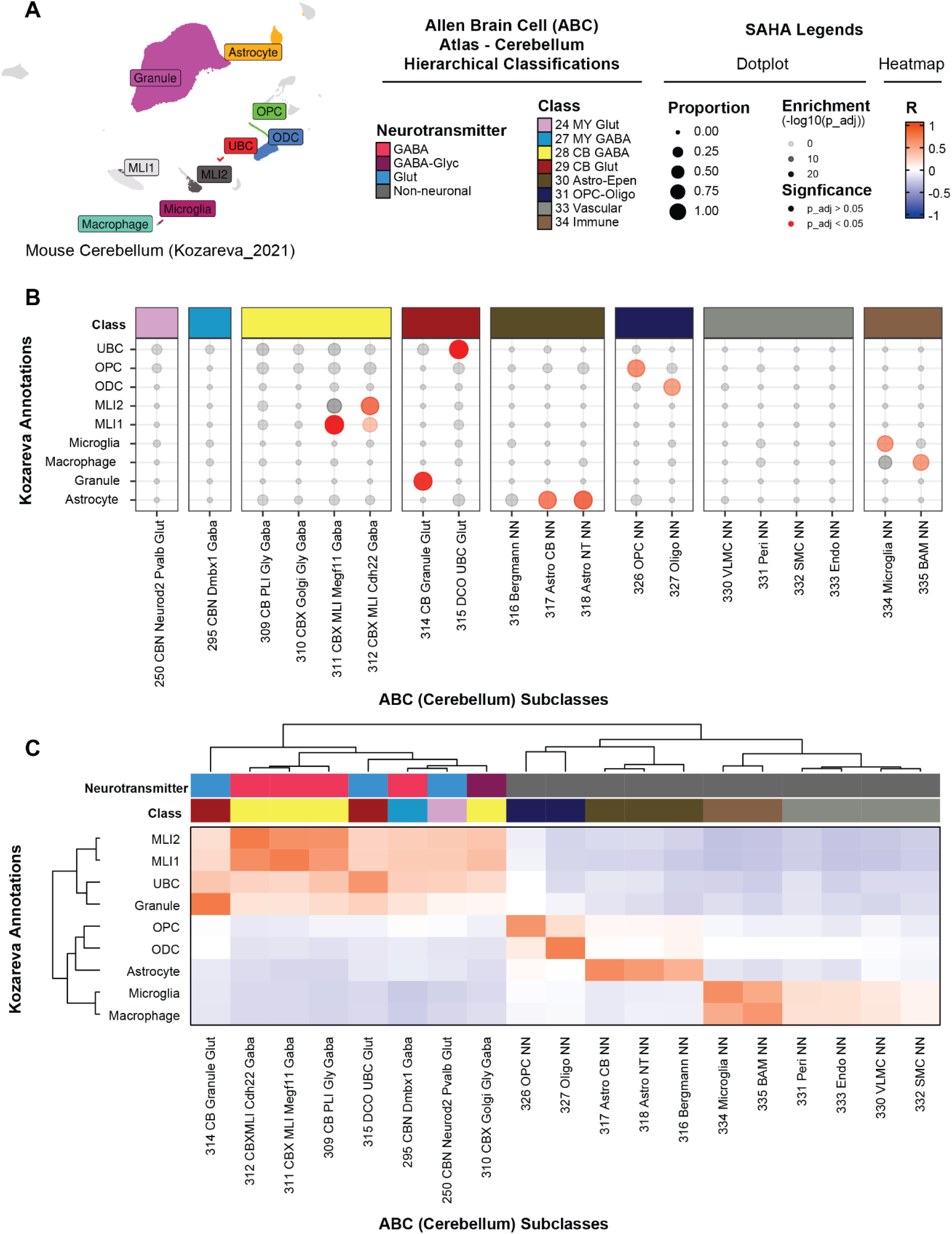
Query-database comparisons leverage similarity scoring for user-based annotation decisions. Cross-study comparison of annotations within the mouse cerebellum reveal high concordance between standard taxonomy (database: ABC Atlas ‘CB’-cerebellum) and study-specific cell annotations (query: 611,034 nuclei object from 6 samples). (**A**) UMAP of publicly available Seurat object colored by annotations found in the object (**See Methods**: “Accession of public datasets”). (**B**) Marker gene similarity between query (y-axis) and database (x-axis) performed via FDR-corrected Fisher’s exact test of positive markers (*SAHA::Run_MarkerBased()*). (**C**) Average gene expression similarity between query (y-axis) and database (x-axis) performed via spearman correlation of variable features from the query dataset (*SAHA::Downsample(…, custom_ds= ‘variable_features_Kozarevà)* against the database (*SAHA::Run_MarkerFree()*). For more information regarding this SAHA run, please refer to the supplement (Table S1-2; Supp Data 1-2).

Notably, 14 clusters resulted in matching ABC subclasses in both marker-based and marker-free analyses that reflected more specifictaxonomic names (i.e. Kozavera_2021 “Golgi” to ABC subclass “310 CBX Golgi Gly Gaba”). Three clusters were inconclusive in marker-based analysis, however offered new interpretations via marker-free analysis (i.e. Kozavera_2021 “Purkinje” to ABC subclass “315 DCO UBC Glut”). Finally, both marker-based and marker-free analyses refined one "Astrocyte" cluster from Kozareva et al. (2021), identifying two distinct ABC subclasses, "317 Astro CB NN" and "318 Astro NT NN," within the broader ABC class "30 Astro-Epen." Taken together, the results of marker-based and marker-free SAHA pipelines reveal a high degree of concordance between the highest matching ABC subclasses and the annotations in the original manuscript.

To demonstrate the utility of SAHA in assigning annotations regardless of the tissue processed, we obtained unprocessed single-cell RNA sequencing of human peripheral blood mononuclear cells (PBMCs) from 10x Genomics demo datasets^33^. As this comparison required a new, non-brain reference dataset, we selected PBMCs from healthy donors in the Covid-19 Multi-omics Blood Atlas^34^ (COMBAT project; 92,205 annotated cells from 124 non-disease donors). SAHA inputs to be used as a reference were calculated using the “Major Subset” annotation column of the processed COMBAT metadata (**See Methods**). After processing the 10x Genomics demo dataset, there were 18 unannotated clusters to be compared to the 17 annotations from the COMBAT project using the SAHA marker-based (**Fig. S2A, Table S3**) and marker-free (**Fig. S2B, Table S4**) pipelines. We re-clustered the data due to apparent overclustering of the demo dataset (**Supp Data 3-4**), resulting in 16 computed clusters that were re-assigned to eight COMBAT project annotations. Notably, four computed clusters were found to best match CD4 cells, four clusters were found to best match monocytes, and two clusters were found to match B cells in the reference (**Fig. S3C-E**). However, computed clusters 8 and 17 did not match any known annotation and were left “Uncertain”. These findings exemplify a common problem in annotation of single-cell data: a single round of clustering can result in over-clustering cells of known biological groupings due to variability present in small clusters of unknown significance.

### Increased refinement of cell type annotations reveals neuronal subclasses without multi-dataset integration

In order to investigate how SAHA can improve the annotation of highly heterogeneous groups of clusters, we obtained a processed, previously published single nuclei RNA sequencing dataset from the cerebral cortex of 4 month old mice^35^. This dataset aimed to describe the effect of inhibiting Inducible Degrader of LDLR (IDOL) in CAMKIIa-expressing neurons of a mouse model of amyloidosis (48,371 nuclei, n = 2 mice per group (5xFAD_IDOL^WT^ vs. 5xFAD_IDOL^Neuron-cKO^)). The original analysis was done with annotations based on computationally defined cluster numbers and broad cell type classification (i.e. cluster 10 inhibitory neurons: “10_iN”). Our goal was to refine cluster labels so that arbitrary naming strategies could be replaced with ABC subclass annotations where applicable.

First, we calculated markers and average expression profiles for clusters at multiple resolutions (0.1 step from 0.1 to 0.9). The number of calculated clusters ranged from 16 clusters at a resolution of 0.1 to 35 clusters at a resolution of 0.9. We then performed both marker-based and marker-free SAHA pipelines iteratively on all resolutions against the ABC reference from the isocortical regions (ISOCTX; **Table S5**). By reporting the consensus best match for every cluster at each resolution, we found many of the subclass annotations to be stable across resolutions (**Fig. S4**). This finding is significant because it means that as higher resolutions create possible over-clustering artifacts, the cells assigned with the final biological term can be preserved without the need for additional user input. However, we did find that a separation between pericytes (331 Peri NN) and endothelial cells (333 Endo NN) was only apparent at resolutions 0.5 and greater. This was also the clustering resolution initially reported in the original study. To avoid unwanted variability due to overclustering and facilitate direct comparison to previous annotations, a resolution of 0.5 was selected to assign final ABC subclass annotations (24 clusters into 24 ABC subclasses across 9 ABC classes; **Fig. S6-7**; **Supp. Data 5-6**).

Next, by directly comparing the clusters and annotations from the original publication^35^ to the SAHA-refined labels, it is evident that the broad cell type labels are consistent with the class-level ABC annotations (**Fig. 2A**). This was especially the case for non-neuronal cells. For example, 100% of cells from the one cluster of astrocytes (6_A) were re-annotated as “319 Astro TE NN” within the “30 Astro-Epen” class. However, the inhibitory and excitatory neuronal clusters were refined at both class and subclass level. Of the 28,124 cells originally annotated as excitatory neurons, 28,084 (99.85%) retained a glutamatergic ABC label across 2 classes and 12 subclasses. Of the 7,272 cells originally annotated as inhibitory neurons, 7,220 (99.28%) retained a GABAergic ABC label across 3 classes and 6 subclasses. To assist with the visualization of these newly discrete cell clusters, we performed subclustering of glutamatergic, GABAergic, and non-neuronal cells while retaining initial SAHA-defined annotations (**Fig. 2B-E**). This additional level of refinement provides improved rationale for resolution selection and applies more generalizable labels without the need for iterative integration-based label transfer. Notably, the two separate clusters that were found to be proportionally responsive to conditional IDOL knockout (10_iN and 22_iN) were both reclassified by SAHA as “056 Chodl Gaba.” This is also consistent with differential expression findings of the two subclusters which showed similar effects to IDOL knockout. Taken together, ABC label transfer using SAHA resulted in a more diverse and specific classification of neuronal subtypes in a manner that was consistent with previous findings.

**Figure 2.**
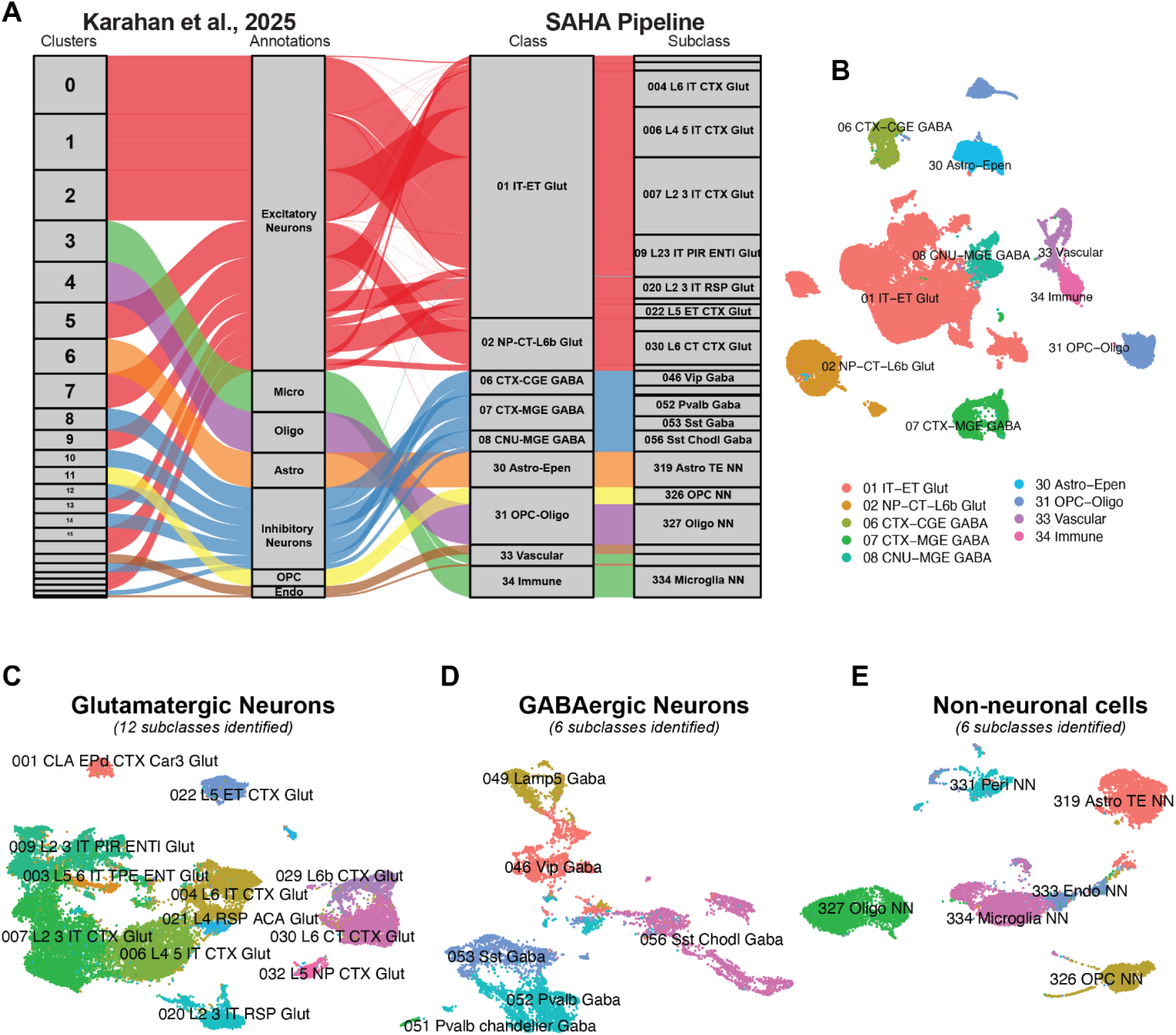
Increased refinement of cell type annotations reveals neuronal subclasses without multi-dataset integration. SAHA-derived annotations using standard taxonomy (database: ABC Atlas ‘ISOCTX’-isocortex) to re-annotate mouse brain cortex snRNA-seq dataset (query: 48,271 nuclei object from 4 samples). Full SAHA outputs are available in the supplemental material (**Fig. S1** ;**See Methods**: “Accession of public datasets”). (**A**) Cell annotation assignment across original study (cluster-level and biologically defined broad cell type) and re-annotation with SAHA (ABC Atlas taxonomy: class and subclass). (**B**) UMAP of publicly available Seurat object colored by SAHA-derived annotations. Subclustering of broad cell groups (ABC Atlas taxonomy: neurotransmitter) following annotation reveal subclasses of (**C**) glutamatergic neurons, (**D**) GABAergic neurons, (**E**) and non-neuronal cell types. Annotations determined by consensus calls between SAHA methodologies (SAHA:AutoAnnotate(…, method = “Both”); **See Methods**: “Interpreting SAHA outputs for cell type determination”). For more information regarding this SAHA run, please refer to the supplement (Table S5-7; Supp Data 5-6).

### Flexible marker-free annotation of spatial transcriptomic assays describe layer-and region-specific cell types

Probe-based targeted transcriptomic assays can increase sensitivity towards genes of interest. However, as probe design is an expensive and resource limiting step of experimental design, the inclusion of cell type specific marker genes is not always possible. To investigate whether the marker-free SAHA pipeline is sufficient to identify layer- and region-specific cell types, we obtained a processed spatial transcriptomic assay from Vizgen MERFISH demo datasets^36^. By further subsetting 9,048 cells within a spatial field of view of the cerebral cortex (**See Methods**), we clustered the cells based on the 481 features present in the assay (**Fig. 3A-B**). Next, we ran the marker-free SAHA pipeline, comparing the 13 region of interest subclusters against the expression profiles of those 481 features within the cell types present in the ABC atlas isocortex and striatum (ISOCTX, STR; **Fig. S5A-B; Table S8-9; Supp Data 7-10**). We report the presence of six non-neuronal cell types, three GABAergic cell types, and four glutamatergic cell types (**Fig. 3C-D**). Notably, the expression profiles of clusters six and seven were negatively correlated to the all non-neuronal cell types but only weakly associated with any neuronal types within the isocortex (**Fig. 3C**). Upon further investigation, the spatial position of cells from these clusters revealed a more medial location than would be expected of cortical neurons. By expanding the investigation to include the ABC striatal region, the identity of clusters six and seven were revealed to be dopaminergic subclasses specific to the midbrain (**Fig. 3D**). By plotting the cortical depth of cells belonging to each of the 7 neuronal and 6 non-neuronal ABC subclasses, cells segregate out by expected layer- and regional-specificity. Layer 2/3, 5, and 6 glutamatergic neurons are arranged in descending cortical depth. Striatal dopaminergic neurons (061 STR D1 Gaba, 062 STR D2 Gaba) are medial to the cortical neurons (**Fig. 3E**). Finally, non-neuronal cells are evenly distributed with the exception of oligodendrocytes (327 Oligo NN) due to the presence of the corpus callosum at a relative cortical depth of approximately 7500-8000 relative assay units (**Fig. 3F**). Taken together, marker-free SAHA analysis of a targeted spatial transcriptomic assay retained the expected organization of all cell types identified.

**Figure 3.**
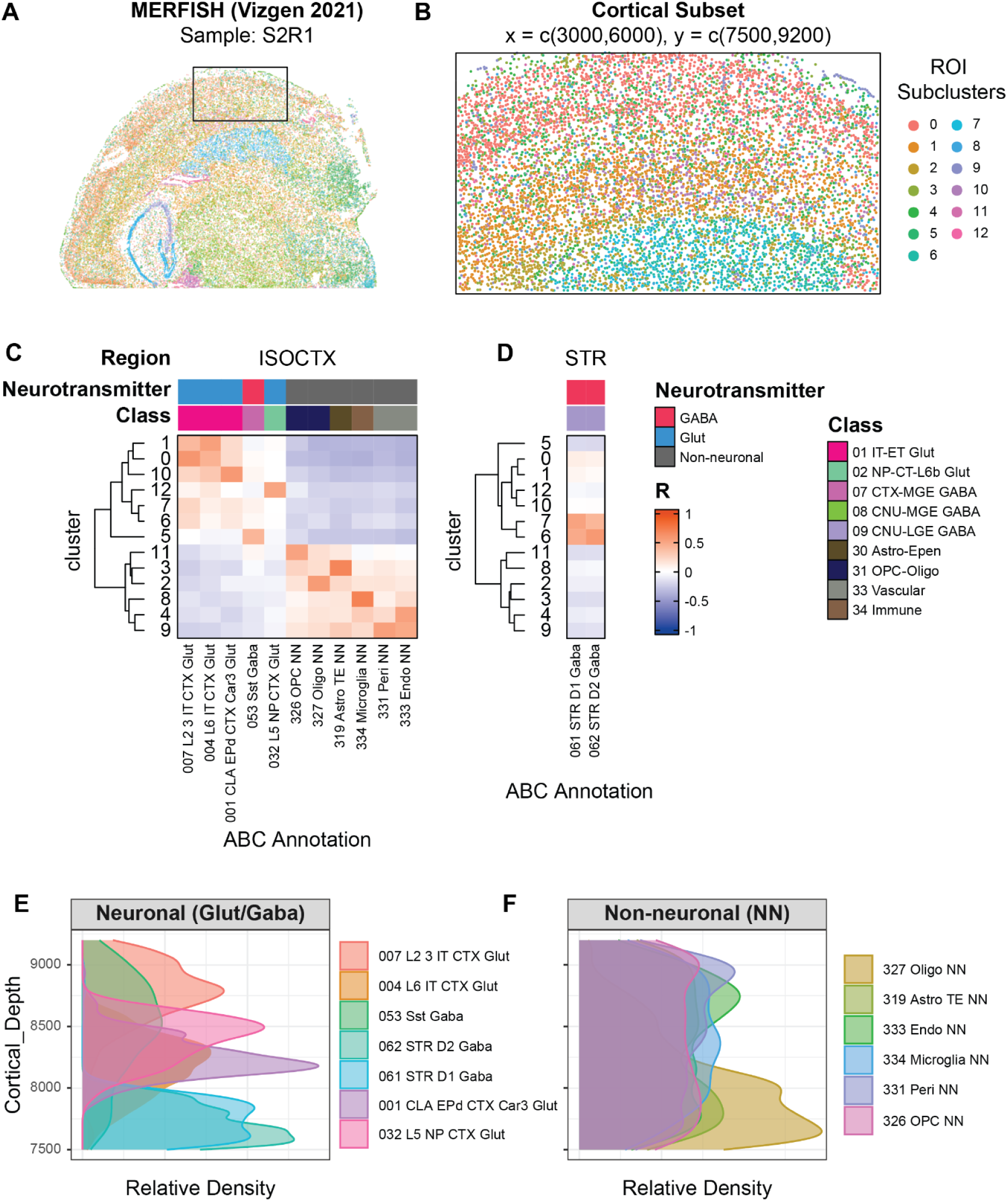
Flexible marker-free annotation of spatial transcriptomic assays describe layer- and region-specific cell types. SAHA-derived annotations using standard taxonomy (database(s): ABC Atlas ‘ISOCTX’-isocortex and ‘STR’-striatum) to Vizgen MERFISH demo dataset. Full SAHA outputs are available in the supplemental material (**Fig. S5** ;**See Methods**: “Accession of public datasets”). (**A**) Region-of-interest (ROI) selected from cortical region of one hemi-coronal section of mouse brain (x = c(3000,6000), y = c(7500, 9200); 8,983 cell object). (**B**) Subclustering of ROI reveals 13 clusters. Marker-free SAHA outputs of (**C**) isocortical and (**D**) striatal cell types. Cortical depth by assay coordinates of (**E**) neuronal and (**F**) non-neuronal cell types. Annotations determined by maximum R^2^ value across both SAHA runs (SAHA:AutoAnnotate(…, method = “AvgExp)). For more information regarding this SAHA runs, please refer to the supplement (Table S8-9; Supp Data 7-10).

### Versatile study-study comparisons reveal human microglia and astrocyte substates across Alzheimer’s disease neuropathology measures

There is a growing body of evidence that certain cell types take on contextual cell states, especially glial cell types during neurodegenerative diseases^37^. One challenge in identifying these states is that they are transient and may not contribute sufficient variability to form discrete clusters^28^. Two strategies to identify these states are to (i) perform cell type-specific subclustering or (ii) identify state-specific gene expression profiles without iterative subclustering. To test whether SAHA is capable of identifying known glial activation states during Alzheimer’s disease (AD), we applied SAHA’s marker-free pipeline to astrocytes and microglia subclustered or pseudobulked by disease group. First, we obtained processed single-nuclei RNA sequencing data from the medial temporal gyrus of AD patients and age matched controls^38^. We used SAHA to compare 44,328 microglia (**Figs. 4A-C, S5A; Table S11; Supp Data 11-12**) and 70,009 astrocytes (**Figs. 4D-F**, **S5B; Table S12; Supp Data 13-14**) to known glial states observed across neurodegenerative diseases^29^.

**Figure 4.**
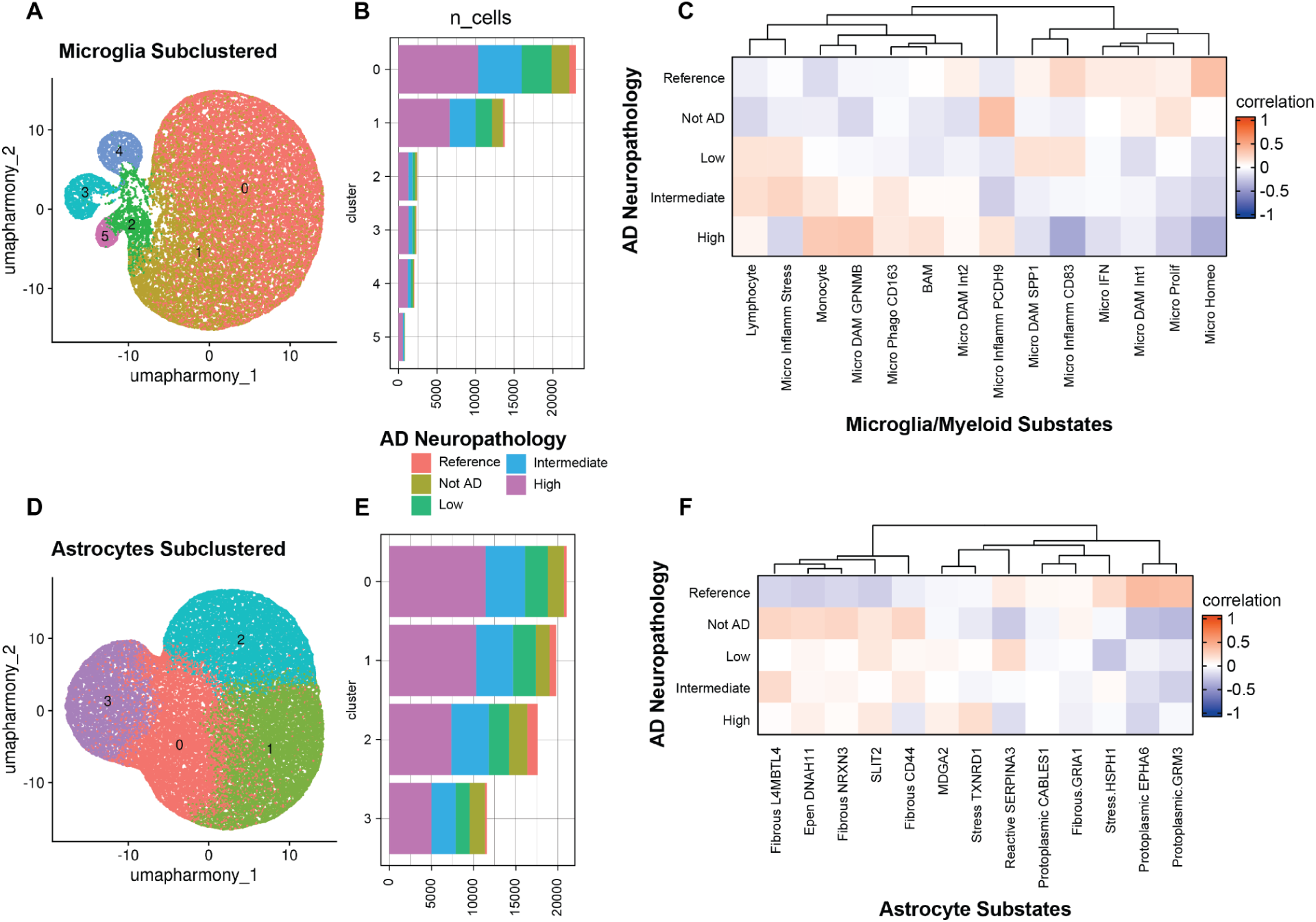
Versatile study-study comparisons reveal human microglia and astrocyte substates across Alzheimer’s disease neuropathology measures. SAHA-derived annotations using custom glial atlases to cells from Seattle Alzheimer Disease (SEAAD) cohort (query: 114,337 cell object from 89 samples). SAHA outputs queried by subcluster are available in the supplemental material (**Fig. S3**; **See Methods**: “Determining level of query for annotation”). (**A-B**) Microglia subclustering reveals 6 clusters across AD neuropathology as reported in the original publication. (**C**) Marker-free SAHA outputs of 14 microglia/myeloid substates. (**D-E**) Astrocyte subclustering reveals 4 clusters across AD neuropathology as reported in the original publication. (**F**) Marker-free SAHA outputs of 13 astrocyte substates (cortical and subcortical substates merged from original publication for comparison). Cross-individual integration and clustering performed with Harmony in Seurat. For more information regarding this SAHA run, please refer to the supplement (Table S10-11; Supp Data 11-14).

Harmony-integrated microglia clustering revealed five subclusters (**Fig. 4A**) evenly spread across reported AD neuropathology groups (**Fig. 4B**). When analyzed at the level of subcluster, the majority of cells (‘0’ = 22,970, ‘1’ = 13,769) appeared to broadly resemble both homeostatic microglia as well as several disease-associated, inflammatory, and phagocytic states (**Fig. S5A**). Other subclusters appeared to better reflect border-associated macrophages (BAMs ;’2’ = 2,425), monocytes (‘5’ = 819), and a PCDH9+ inflammatory states (‘3’ = 2,301, ‘4’ = 2,044). The apparent lack of clear homeostatic or disease-associated subclusters is not surprising given that AD neuropathology groups were evenly represented across the integrated subclusters. The ambiguity between clusters and cells attributable to disease conditions is a downside of the aforementioned strategy to identify cellular states via clustering, where integration can possibly ablate disease-relevant clustering variation. However, when average expression profiles were calculated by pseudobulking all microglia across AD neuropathology groups rather than subclusters, SAHA reveals the expected annotation patterns (**Fig 4C**). Namely, reference microglia were homeostatic, age-matched non-pathological controls (‘Not AD’) were inflammatory but lacked similarity to disease-associated states, and increasing burden of AD neuropathology resulted in the appearance of a GPNMB+ DAM state. These findings emphasize the sensitivity of state-identification to decisions regarding integration and data partitioning, however SAHA offers the advantage of enabling the user to make rapid comparisons to previously published states without the reliance on marker genes.

Harmony-integrated astrocyte clustering revealed four subclusters (**Fig. 4D**) evenly spread across reported AD neuropathology groups (**Fig. 4E**). While less is known regarding astrocyte states in response to disease, the major divide between protoplasmic- and fibrous-states is well described^39^. When analyzed at the level of subcluster, cells delineated across the protoplamismic-fibrous spectrum (**Fig. S5B**) with one clear stress-associated astrocytic subcluster (‘0’ = 21,028). When average expression profiles were calculated by pseudobulking all astrocytes across AD neuropathology groups, we found that the reference group was a major contributor to protoplasmic astrocyte states while AD-patients and age-matched controls were largely fibrous. The stress-associated state (Stress TXNRD1+) mapped to a subcluster that was neither strongly fibrous nor protoplasmic and was enriched in high AD neuropathology. This pattern suggests the presence of a disease-associated astrocyte state that has not yet been fully characterized. Similar to the characterizations of microglia, these findings demonstrate how SAHA may enable researchers to rapidly evaluate astrocyte identity without the reliance on marker genes.

## DISCUSSION

Annotating cell types or cell states from single-cell RNA sequencing data is the rate-limiting step which determines statistical power, biological resolution, and utility of any dataset. In particular, datasets from the mammalian brain are highly heterogeneous and contain transient, context-specific cell states across development^40^, non-pathological aging^41–43^, and disease^23,29,44–46^. Neuronal diversity may be defined by classes varying in neurotransmitter release^47^, circuit and layer projection^48^, or susceptibility to disease^49^. On the other hand, glial diversity is most commonly associated with regional heterogeneity^50^ or polarization during stimuli^51^. Once these states are identified in a newly collected dataset, the number of overall cells or donors contributing to a given cluster informs what downstream analyses are appropriate. Although no gold standard exists for determining power or standard usage of single-cell data, innovative new technologies are beginning to enable integration of epigenetic and spatial data^52^. These increasingly rich datasets compound the problem of cell type annotation, increasing the dimensionality of multi-omic features and therefore computational cost of integration within and across studies.

For studies of the mammalian brain, large consortia have created consensus labeling strategies with a special focus on neuronal heterogeneity^20^. These studies have enabled discovery of subtype-specific alterations in studies of neurodegenerative diseases^23,44–46^, schizophrenia^53^, and other neurological disorders^54^. Reproducibility challenges remain where independent sample series omit canonically present cell types or create study-specific labels that do not generalize. Secondary meta-analysis has enabled some cross-study comparisons^29,55^, however the largest challenge of determining a taxonomy for a single-cell dataset is inherently a problem of cluster annotation.

The two major approaches of cell type annotation are through label transfer following integration or *de novo* label creation based on biological investigation. The latter we refer to as “hand annotation”, as it involves investigating the expression and specificity of individual genes in order to create a new label specific to a study. However, integration-based annotation has become increasingly popular with computational biologists for its reproducibility and transparency. Iterative annotation and cleaning using both approaches poses several challenges: i) the possibility of poorly predicted clusters, ii) deterministic label transfer improperly matched between query and reference, and iii) the necessity to fully integrate query and reference. Several command-line^56,57^ and web tools^58,59^ are available to perform integration-based clustering, however the memory and bandwidth constraints remain major limiting factors. We propose that summary-level statistics of marker genes and average expression profiles offer a scale-free alternative to annotation with either deterministic automation or step-wise semi-automated biological investigation. This approach which we call “semi-automated hand annotation” (SAHA) offers rapid and easily reportable annotation which can be applied across datasets using minimal computational resources.

SAHA implements a Fisher’s exact test^60^ for per-cluster markers and a correlation^61^ of scaled, normalized average expression profiles to aid in annotation. The marker-based strategy closely resembles traditional hand annotation implemented since the commercialization of single-cell assays^62,63^. However, the marker-free strategy more closely resembles post hoc analyses to demonstrate similarity across species or models^22^. Both approaches apply a reportable statistic that can be used to justify annotation. We found that applying this strategy to an annotated mouse cerebellar dataset^32^ (Kozareva_2021) and the cerebellar cells from the ABC atlas^20^ appropriately matched cell type annotations (**Figure 1**). Aside from confirming the fidelity of the original annotation, SAHA also provides additional information regarding which genes drive the similarity across studies and similarity across cell types. For example, astrocytes from Kozareva_2021 contained significant overlap with cerebellar astrocytes (“317 Astro CB NN”), nontelencephalic astrocytes (“318 Astro NT NN”), and Bergmann glia (“316 Bergmann NN”) in the ABC atlas. The SAHA outputs enable annotation with a low barrier to entry but are also useful to investigate cell types identities.

Many studies adopt a combination of broad cell type labels with study-specific cluster numbers or markers (i.e. InN_1 or InN_GAD1). Using the region-specific taxonomic designations for neurons in the ABC atlas, we demonstrated how SAHA can offer more reproducible labels to study-specific annotations (**Figure 2**). With the addition of ABC annotations, the two clusters which were depleted in a mouse model of amyloidosis^35^ with the deletion of an E3 ubiquitin ligase (IDOL) (clusters 10 and 22) were both annotated as Somatostatin positive chondrolectin expressing inhibitory neurons (056 Sst Chodl Gaba), a distinct subtype of neurons known to project from the cortex to the hippocampus and hypothalamus^64^. In this case, post-hoc annotation of cell types confirms that these similarly affected cell types are likely overclustered cells of similar function. Interestingly, the Sst_Chodl subtype comprises long-range GABAergic projecting neurons that mature late in development and have been shown to also be selectively vulnerable to neuropsychiatric risk associated with sleep disturbances^65^. The original finding that subtypes of inhibitory neurons are uniquely sensitive to conditional knock out of IDOL is therefore bolstered by the ability to further classify their neuronal identity. This sort of refinement may be done during initial annotation of new sample series or as a meta-analysis of published findings to help biologists draw conclusions from hand-annotated omic-heavy datasets.

Beyond the application of annotating clusters from single-cell/nuclei assays, rapid annotation also enables optimization of cluster resolution and investigation of cell identities that are poorly labeled by the cell types present in the corresponding atlas. First, we showed that applying to clustering across resolutions revealed divergence of biologically distinct clusters as a rationale for selecting a resolution (**Fig. S3**). Second, we showed how sampling contamination of non-cortical neuronal subtypes could be correctly identified as dopaminergic neurons medial to the corpus callosum (**Fig. 3**). In some cases, this flexibility may aid in the identification of less common or rare cell types instead of forcing the highest probability annotation from an integrated reference. Finally, with a defined set of microglia and astrocyte substates, we were able to attribute annotations to mathematically defined clusters (**Fig. S5**) and pathologically scored disease groups (**Fig. 4**). These further applications of SAHA reveal the advantage of non-deterministic, rapid discovery of cell type identities. Finally, all decisions in this pipeline are reported via html reports (see **Supp. Data**) and easily sharable as “SAHA objects” that can be saved and shared like any other processed single-cell data.

The distinct advantage of this approach is that defining markers and identity remains firmly in the hands of the user. By enabling a “semi-automated” decision making process, we ensure that the results are driven by biological expertise rather than tuned computational models. Furthermore, this workflow does not employ LLMs or machine-learning strategies that require the user to access and input additional datasets that may be storage-restrictive or held behind data use agreements. By lowering the hardware and coding barriers, we enable usage by novice coders and direct investigation by biological experts. Future work will focus on applying similar strategies to non-gene markers of cell types, such as single-cell protein and epigenetic measures. While species-specific databases and availability of summary-level metadata are current limitations of SAHA implementation, consortia of laboratories with single-cell data who commit to making marker and expression data publicly available will address a critical need of the single-cell community. Finally, benchmarking hand annotation to the many integration-based and machine-learning enabled annotation strategies will be necessary to determine a gold standard for the community.

## MATERIALS AND METHODS

### Public dataset processing

Public datasets were accessed for two purposes: i) use as query samples to attempt annotations^32,33,35,36,38^ and ii) to use as database references^20,29,30^ for label transfer via a summary statistic-based, non-integration method. Databases were generally used without re-processing, additional clustering, or further filtering. To this end, annotations at the reported level were used to calculate marker genes (*Seurat::FindAllMarkers()*) and calculate average expression profiles (*Seurat::AverageExpression()*). For datasets that exceed reasonable matrix constraints in R, marker genes and average expression profiles were calculated in Python.

Annotations from three studies were used to create database reference inputs in this study: all cell types from multiple regions of the mouse brain^20^, human glial substates derived from integrative analysis of 4 multi-region neurodegeneration single nuclei atlases^29^, and PBMCs from individuals infected with influenza, COVID-19, or healthy controls^30^. From the mouse ABC atlas^20^, processed datasets were obtained through the ABC Atlas API (“Whole mouse brain clustering”; version 20231215; Accessed July 17, 2024). Cell types present in multiple regions (i.e. 338 in 12 regions; 171 region-specific, 5 shared in all regions, and 162 shared between 2-11 regions) were recalculated with each appearance as marker gene calculation and downstream expression normalization are respective to the other cell types present. Average expression profiles and markers from the mouse ABC atlas were integrated into SAHA functionality by deposition into the *SAHAdata* package^31^. For a human brain glia reference, samples from four atlases were processed as described^23,44–46^. In brief, raw sequencing data were obtained under DUA and reprocessed through a doublet- and ambient-RNA- conservative pre-processing pipeline. Astrocytes and microglia from these studies were integrated and annotated using consensus glial substate marker sets published between 2017 and 2024. For a human PBMC reference, the COMBAT2022 study was accessed via CombatDB online server and labels at the reported “major_subset” level were used for the calculation of markers and average expression profiles. Input files for studies not included in the *SAHAdata* package are included in the supplement of this study.

Transcriptomic datasets from five sources were used to create query inputs from this study: a mouse cerebellum single-cell study published prior to the ABC atlas^32^, a mouse snRNAseq dataset profiling the isocortex without celltype enrichment or sorting^35^, spatial transcriptomic data available from Vizgen^36^, glial cells from the medial temporal gyrus of the SEA-AD cohort^38^, and PBMC data available from 10x Genomics^33^. For the mouse cerebellum dataset, markers and average expression profiles were created from reported annotations to confirm fidelity to known cell types in the ABC atlas. For the mouse isocortex dataset, markers and average expression profiles were created from annotations concatenating broad cell type and cluster number (i.e. ExN_1, Oligo_2, etc.). For the spatial transcriptomic dataset, one sample containing a single hemisphere of the mouse brain was subset down to coordinates containing the cerebral cortex. Clustering was performed in *Seurat* to identify region-of-interest-specific cell clusters and used for the creation of summary statistic inputs. The SEA-AD cohort samples were downloaded with annotations as reported. The 70,009 astrocytes and 44,328 microglia were subset in Python then transferred to Seurat using the package *schard*^66^. Sub-clustering was performed separately for each glial cell type using *Seurat*, and summary statistics were calculated at the levels of cluster and of “AD Neuropathology Group”, as reported in the sample metadata. Finally, h5 files for human PBMC were downloaded from the 10x Genomics dataset^33^ “PBMCs from a healthy donor: Whole transcriptome analysis”, processed in Seurat, and cluster-level input files were created. For specifics regarding principal component selection, Louvian clustering resolution, integration, and subsetting, please refer to the code available on GitHub (https://github.com/jungsukimlab/SAHA_manuscript) and Zenodo1.

### Preparing the SAHA pipeline inputs

The preparation of SAHA inputs rely on either marker gene calculation for marker-based analysis or average expression profile calculation for marker-free analysis. Marker-based analysis was implemented with *Seurat::FindAllMarkers(....,only.pos=T, logFC > 0.25)* calculating per cluster markers with a Wilcoxon rank sum test. SAHA allows any method for calculation of markers, however recognizes dataframes where row names are genes and column headers include: p_val, avg_log2FC, pct.1, pct.2, p_val_adj, and cluster. Marker-free analysis performed with *Seurat::AverageExpression(layer=”data”)* calculates the average of each scaled and log-normalized value per gene within a given grouping of annotations. SAHA also allows for any calculation of average expression, however recognizes dataframes where row names are genes and column headers are the names cluster or annotation groupings. For data preparation outside of Seurat, markers and average expression were calculated using comparable statistical decisions and file outputs were renamed to align with those of *Seurat::FindAllMarkers()* or *Seurat::AverageExpression()*. Finally, Marker-free analysis contains an optional down-sampling parameter which necessitates the storage of the top 2,000 variable features used in the database reference clustering. Variable features were calculated with *Seurat::FindVariableFeatures()* or corresponding functions outside of Seurat. To prepare all SAHA pipeline inputs from a single Seurat object, the function *SAHA::Seurat2SAHA()* implements the preparation of markers, average expression, and variable features as described.

### Marker-based annotation analysis

Marker-based analysis aimed to compare defined marker sets from the query to the database. Initial SAHA object creation checked for feature concordance as a confirmation that feature descriptors were common between the two datasets (i.e. ENSMBL ID’s, gene names, etc). Automatic filtering was performed on both query and database marker sets for significance (p < 0.05), effect (FC > 1.5), specificity (pct.2 < 0.25, or less than 25% of off-target cells containing a count), and sensitivity (pct.1 > 0.75, or more than 75% of on-target cells containing a count) per cluster. Additional tuning was performed via *SAHA::TuneMarkers()* to select top 100 markers per cluster in the database marker set. Over-representation analysis of enriched markers was performed via hypergeometric test as implemented in *stats::dhyper()* accounting for size of query cluster set, size of database cluster set, overlap, and all genes used as markers at least once across all comparisons. Significant enrichment of cluster markers in a database annotation was reported as the p-value of the hypergeometric test and the proportion of overlap, FDR corrected across all comparisons. Further tuning and parameter setting is specified per SAHA run in the code available on (https://github.com/jungsukimlab/SAHA_manuscript).

### Marker-free annotation analysis

Marker-free analysis aimed to compare the expression of all, or a specified downsampled subset of, genes between clusters of the query-set versus annotations in the database. In preparation for drawing a correlation between genes in each cluster x annotation combination, we first confirmed that feature descriptors were common between the two datasets (i.e. ENSMBL ID’s, gene names, etc). Next, the average expression of each gene was z-scored across each cluster or annotation in the respective feature set–this default setting (“across-cluster”) was implemented to maximize the relative utility of each gene across same-dataset groupings. Optionally, users could instead apply the “within-cluster” setting, in which all genes are z-scored within each individual cluster or annotation. Correlation analysis was then performed using base R. While the correlation test defaults to using Pearson’s correlation, there is flexibility to include Spearman or other specified methods within *base::cor()*. Optional down-sampling to custom gene sets or the variable features from the query dataset specified per SAHA run can be found in the code available on (https://github.com/jungsukimlab/SAHA_manuscript).

### Interpreting SAHA outputs for cell type determination

Interactive cluster annotation was performed with *SAHA::SemiAutoAnnotate()* to visualize each query cluster in RStudio or via *SAHA::AutoAnnotate()* matching the best annotation for marker-based (minimum p-value across cluster x annotation combination) or marker-free (maximum correlation coefficient across cluster x annotation combination). Further refinement enabled by biological insights specified per SAHA run can be found in the code available on GitHub (https://github.com/jungsukimlab/SAHA_manuscript). Notably, SAHA runs were iterated over group subsets, resolution definitions, and regional references to improve annotation refinement.

## Abbreviations

ABC: Allen Brain Cell Atlas
CB: Cerebellum (ABC abbreviation)
COMBAT: Covid-19 Multi-omics Blood Atlas
ISOCTX: Isocortex (ABC abbreviation)
MERFISH: Multiplexed Error-Robust Fluorescence in Situ Hybridization
PBMC: Peripheral Blood Mononuclear Cells
SAHA: Semi-Automated Hand Annotation
SEA-AD: Seattle Alzheimer’s Disease Brain Cell Atlas
STR: Striatum (ABC abbreviation)

## Supplemental data

Supplemental data include markers, average expression, html summaries, and SAHA ‘ann’ objects for mouse cerebellum (Table S1-2, Supp Data 1-2), human PBMC (Table S3-4, Supp Data 3-4), mouse isocortex (Table S5-7, Supp Data 5-6), mouse spatial cortex (Table S8-9, Supp Data 7-10), human microglia (Table S10, Supp Data 11-12), human astrocytes (Table S11, Supp Data 13-14), and a source manifest (Table S12). Supplemental figures include results from mouse cerebellum (Fig. S1), human PBMC (Fig. S2), resolutions across mouse isocortex (Fig. S3), mouse spatial cortex versus isocortical and striatal references (Fig. S4), and glia compared to neurodegenerative substate annotations (Fig. S5).

## DECLARATIONS

### Ethics approval and consent to participate

Not applicable.

### Consent for publication

Not applicable.

### Availability of data and materials

The R package ‘SAHÀ is freely available under MIT licence at https://www.github.com/neurogenetics/SAHA. The code necessary to reproduce figures associated with this manuscript are available at https://www.github.com/jungsukimlab/SAHAmanuscript. Data for accessing the ABC Atlas summary-level statistics used for the SAHA pipeline is freely available under MIT licence at https://www.github.com/neurogenetics/SAHAdata. No single-cell and spatial datasets were created in this study. Information regarding their accession can be found in methods (“Accession of public datasets”).

### Authors’ Contributions

D.J.A., M.R.C., and J.K. conceptualized the study and planned experiments. Software development was performed by D.J.A., R.M., L.H-P., M.E.M., and J.R.G.. Investigation and formal analysis was performed by D.J.A., L.H-P., L.C.D., J.H.P., K.H., H.K.. Funding of this study was obtained by M.R.C. and J.K. and the first draft of the manuscript was written by D.J.A.. All authors reviewed and edited the manuscript. M.R.C. and J.K.. supervised the work.

## Acknowledgements

We would like to thank Xylena Reed, B. Adam Catching, and Cory Weller for their feedback and thoughtful discussion concerning single-cell annotation pipeline development. We would like to especially thank the patients, patient families, and physicians who contributed to the human data included in this manuscript. Information regarding publicly available datasets or those with data use agreements from the Gene Expression Omnibus or Seattle-AD (SEA-AD) consortia are available in the Supplementary Information (Table S12). Full acknowledgement of respective studies, brain banks, and data use agreements available in the Supplementary Information (Supp Data).

## Ethics and Generative AI Use Statement

Large language models (LLMs) were employed for creating documentation of the SAHA pipeline and for annotating code implementing SAHA, practices within current guidance of the NIH (https://osp.od.nih.gov/policies/artificial-intelligence/). We, the authors, affirm that no LLMs were used in the study design, drafting of the manuscript or analyzing any data associated with this study.

## SUPP FIGURES

**Figure S1.**
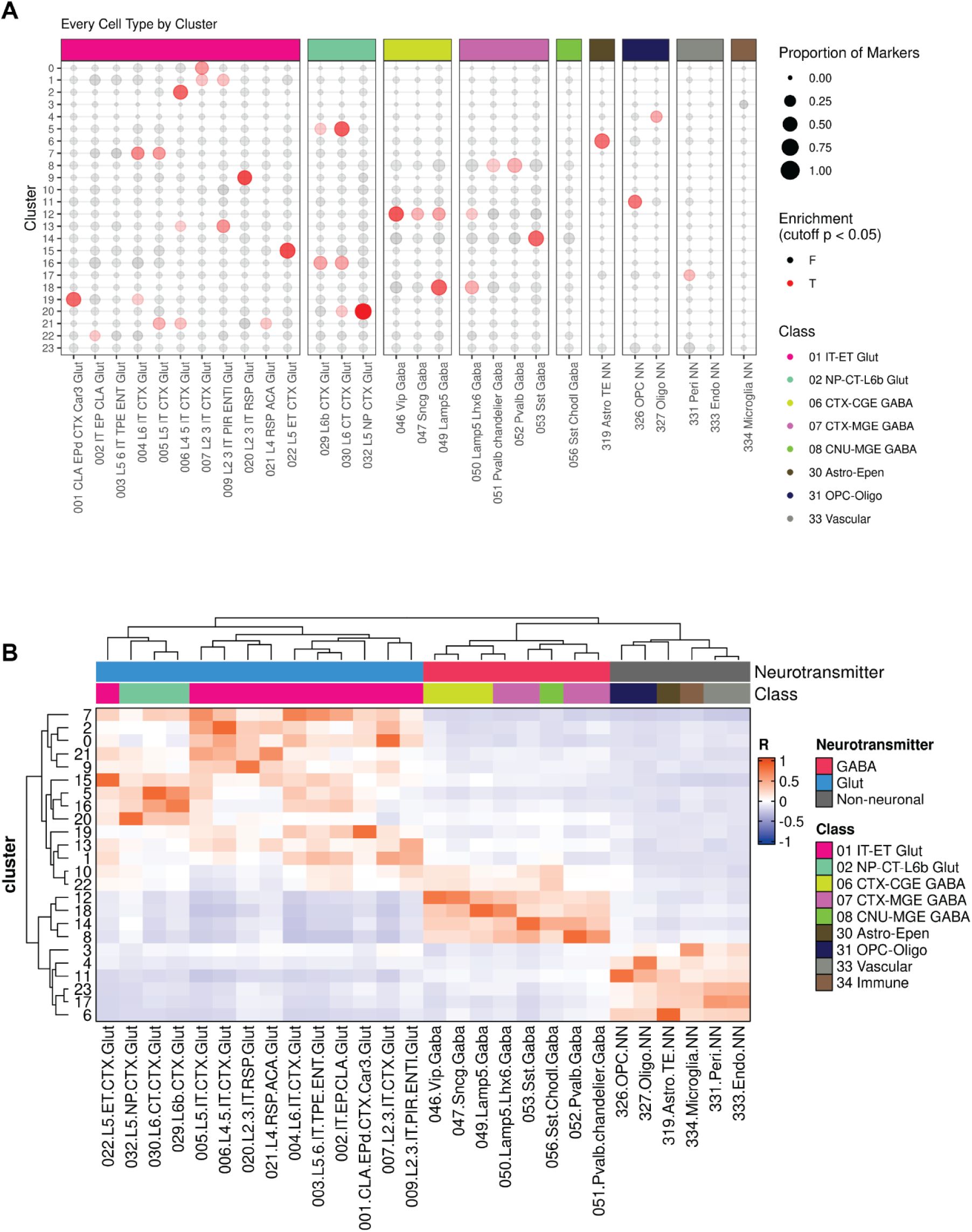
Query-database comparisons leverage similarity scoring for user-based annotation of cortical mouse brain snRNAseq dataset. Cross-study discovery of annotation for clusters within the mouse isocortex reveal high concordance using standard taxonomy (database: ABC Atlas ‘ISOCTX’-isocortex) to re-annotate mouse brain cortex snRNA-seq dataset (query: 611,034 cell object from 6 samples). (**A**) Marker gene similarity between query (y-axis) and database (x-axis) performed via fdr-corrected Fisher’s exact test of positive markers (*SAHA::Run_MarkerBased()*). (**B**) Average gene expression similarity between query (y-axis) and database (x-axis) performed via spearman correlation of variable features from the query dataset (*SAHA::Downsample(…, custom_ds= ‘variable_features_Karahan’)* against the database (*SAHA::Run_MarkerFree()*). Annotations determined by consensus calls between SAHA methodologies (*SAHA:AutoAnnotate(…, method = “Both)*; **See Methods**: “Interpreting SAHA outputs for cell type determination”).

**Figure S2.**
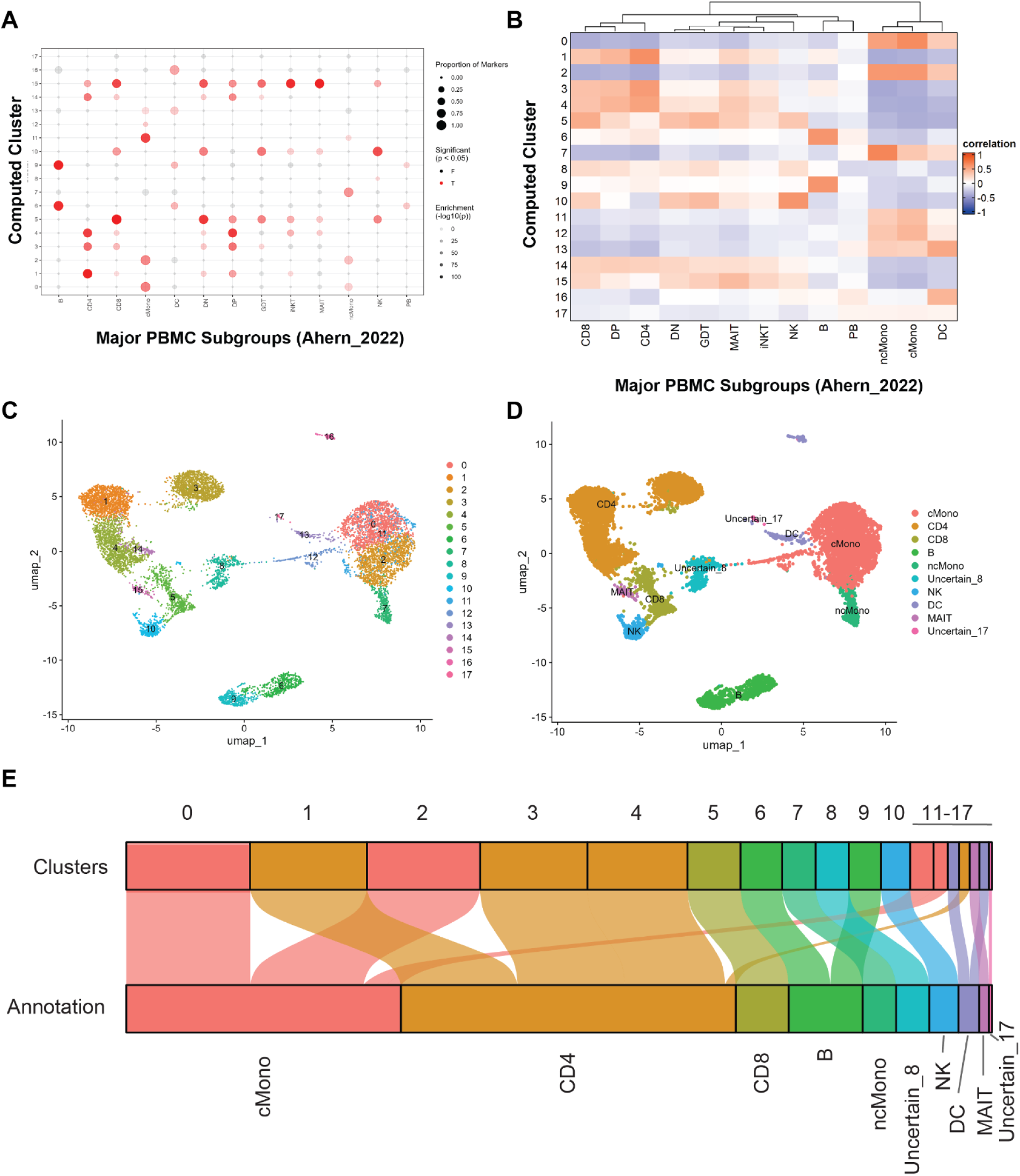
Query-database comparisons leverage similarity scoring for user-based annotation of 10xGenomics PBMC demo dataset. Cross-study discovery of annotation for of human PBMCs (query: 10,204 cell object from one sample^33^) using an annotated cell atlas of healthy PBMCs (database: COMBAT2022 healthy PBMCs (836,148 cells from 124 samples). (**A**) Marker gene similarity between query (y-axis) and database (x-axis) performed via FDR-corrected Fisher’s exact test of positive markers (*SAHA::Run_MarkerBased()*). (**B**) Average gene expression similarity between query (y-axis) and database (x-axis) performed via spearman correlation of variable features from the query dataset (*SAHA::Downsample(…, custom_ds= ‘variable_features_10xPBMC’)* against the database (*SAHA::Run_MarkerFree()*). UMAP of publicly available Seurat object colored by (**C**) clustering and (**D**) annotation (**See Methods**: “Accession of public datasets”). (**E**) Cell annotation assignment across initial clustering (cluster) and annotation with SAHA (COMBAT2022 broad annotations: “Major Type”). Annotations determined by consensus calls between SAHA methodologies (SAHA:AutoAnnotate(…, method = “Both); **See Methods**: “Interpreting SAHA outputs for cell type determination”).

**Figure S3.**
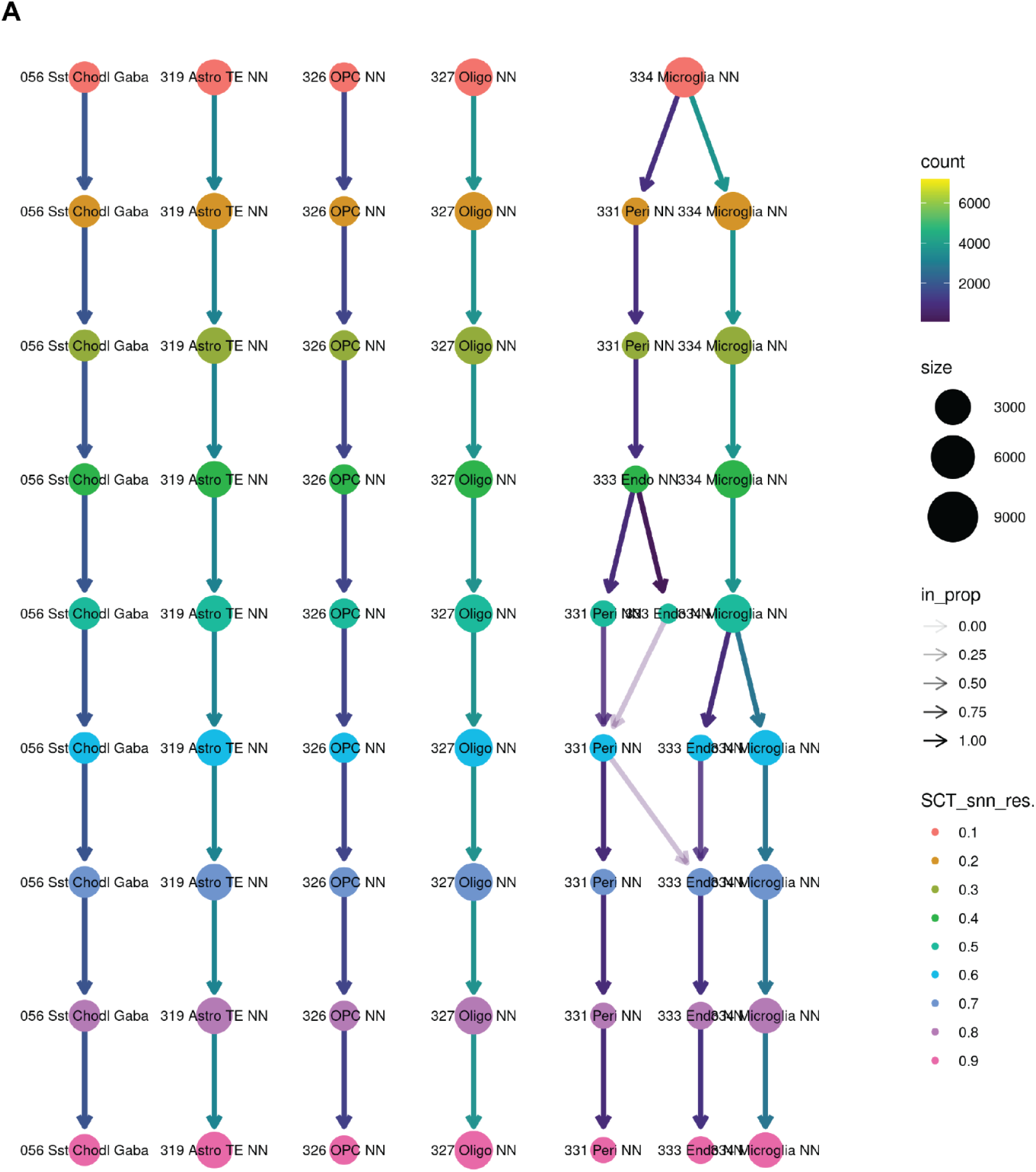
Iterative SAHA runs across multiple queries enables fast biological description for resolution selection. Clustree^67^ of cells grouped by SAHA-derived annotation of mouse brain cortex at nine different clustering resolutions (Figure 2, **Figure S1**). Annotations determined by consensus calls between SAHA methodologies (SAHA:AutoAnnotate(…, method = “Both)).

**Figure S4.**
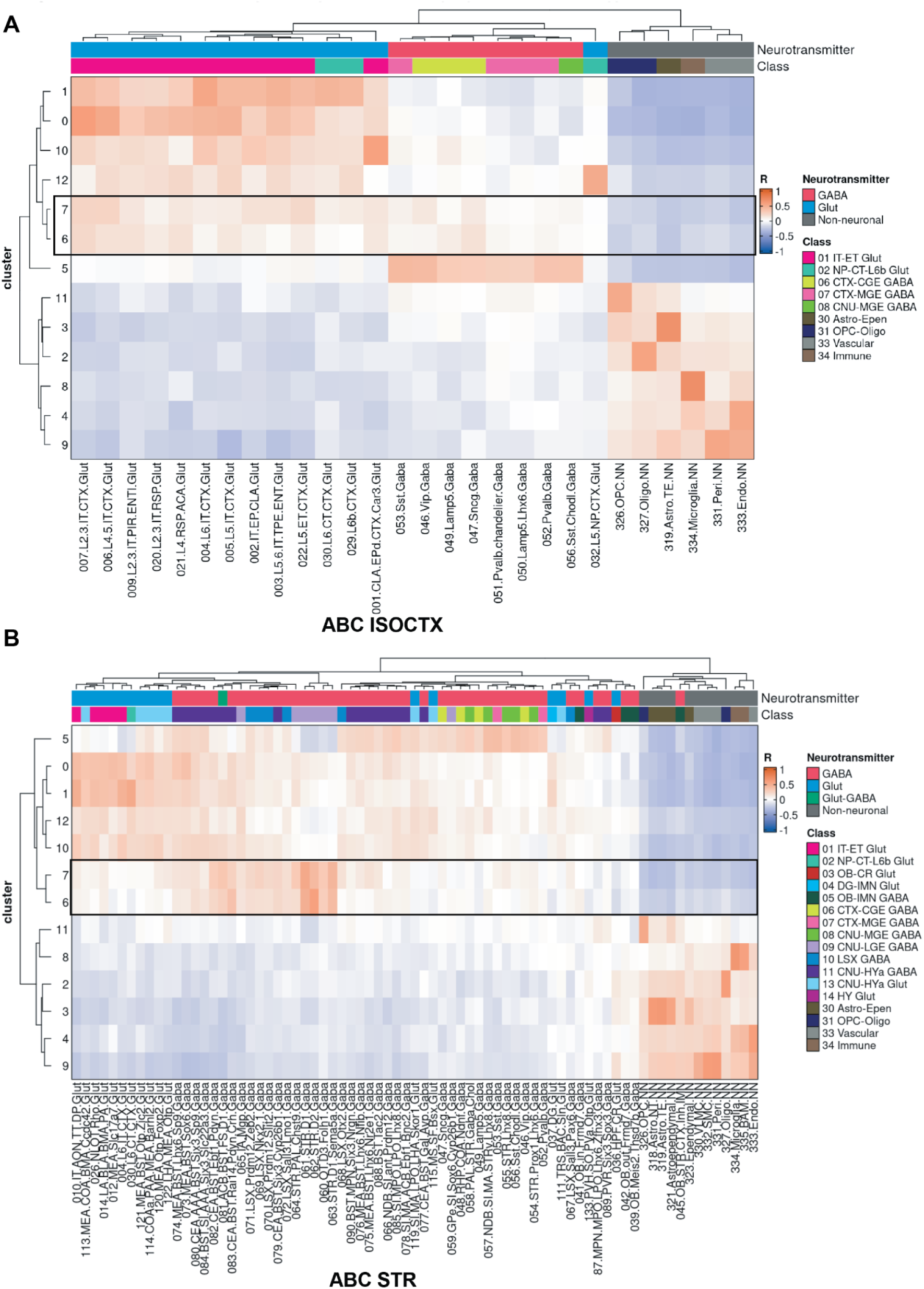
Iterative SAHA runs across multiple databases enables fast biological description of regionally-defined cell types. SAHA-derived annotations using standard taxonomy (database(s): ABC Atlas (**A**) ‘ISOCTX’-isocortex and (**B**) ‘STR’-striatum) to Vizgen MERFISH demo dataset^36^. Average gene expression similarity between query (y-axis) and database (x-axis) performed via spearman correlation of variable features from the query dataset (*SAHA::Downsample(…, custom_ds= ‘variable_features ‘)* against the database (*SAHA::Run_MarkerFree()*). Annotations determined by maximum R^2^ value across both SAHA runs (*SAHA:AutoAnnotate(…, method = “AvgExp)*). Clusters 6 and 7 (black box) were determined to be striatal neurons while other types appropriately matched isocortical cell types.

**Figure S5.**
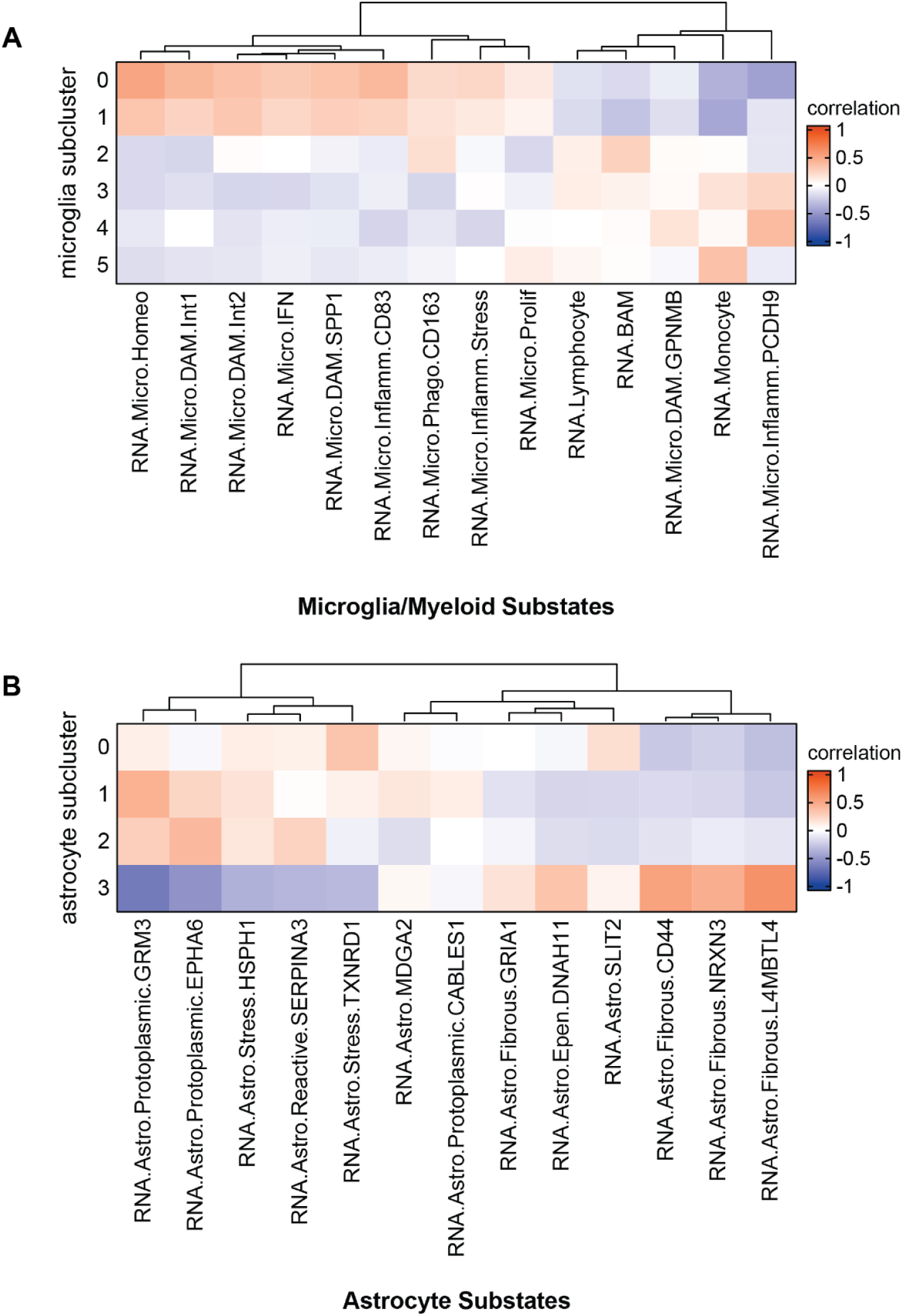
Query-database comparisons leverage similarity scoring for user-based annotation of human microglia and astrocyte snRNAseq dataset. Cross-study comparison of annotations using custom glial atlases to (**A**) microglia and (**B**) astrocytes from SEAAD cohort (query: 114,337 cell object from 89 samples). Average gene expression similarity between query (y-axis) and database (x-axis) performed via spearman correlation of variable features from the query dataset (*SAHA::Downsample(…, custom_ds= ‘variable_features_SEAAD’)* against the database (*SAHA::Run_MarkerFree()*).

## Supplemental Data

Guide to supplemental data available on Zenodo (https://doi.org/10.5281/zenodo.20691702)^68^.

Supplemental Table 1. **Marker gene expression results for mouse cerebellum.** Data from published dataset<sup>32</sup> against annotation from ABC mouse brain atlas.
| Supp Data | Description | Associated Figure |
| --- | --- | --- |
| 1 | cerebellum report html | Fig. 1 |
| 2 | cerebellum completed ann | Fig. 1 |
| 3 | PBMC report html | Fig. S2 |
| 4 | PBMC completed ann | Fig. S2 |
| 5 | mouse sn report html | Fig. 2 |
| 6 | mouse sn completed ann | Fig. 2 |
| 7 | mouse spatial isoctx report html | Fig. 3 |
| 8 | mouse spatial isoctx completed ann | Fig. 3 |
| 9 | mouse spatial str report html | Fig. 3 |
| 10 | mouse spatial str completed ann | Fig. 3 |
| 11 | micro html | Fig. 4 |
| 12 | micro ann | Fig. 4 |
| 13 | astro html | Fig. 4 |
| 14 | astro ann | Fig. 4 |

Supplemental Table 1. **Marker gene expression results for mouse cerebellum.** Data from published dataset^32^ against annotation from ABC mouse brain atlas.

Supplemental Table 2. **Average expression results for mouse cerebellum.** Data from published dataset^32^ against annotation from ABC mouse brain atlas.

Supplemental Table 3. **Marker gene expression results for human PBMC.** Data from published dataset^33^ against annotation from COMBAT PBMC atlas.

Supplemental Table 4. **Average expression results for human PBMC.** Data from published dataset^33^ against annotation from COMBAT PBMC atlas.

Supplemental Table 5. **Cell taxonomy including multiple annotations for resolution selection.** Data from published dataset^35^ against annotation from ABC mouse brain atlas.

Supplemental Table 6. **Marker gene expression results for mouse isocortex.** Data from published dataset^35^ against annotation from ABC mouse brain atlas.

Supplemental Table 7. **Average expression results for mouse isocortex.** Data from published dataset^35^ against annotation from ABC mouse brain atlas.

Supplemental Table 8. **Average expression results for mouse spatial transcriptomics versus reference isocortex.** Data from published dataset^36^ against annotation from ABC mouse brain atlas.

Supplemental Table 9. **Average expression results for mouse spatial transcriptomics versus reference striatum.** Data from published dataset^36^ against annotation from ABC mouse brain atlas.

Supplemental Table 10. **Average expression results for human microglial substates.** Data from published dataset^38^ against annotation from integrated glial atlas.

Supplemental Table 11. **Average expression results for human astrocytic substates.** Data from published dataset^38^ against annotation from integrated glial atlas.

Supplemental Table 12. **Data source manifest.** Accession numbers and data location for each public dataset utilized in this study.

Supplemental Data 1. **HTML report expression of mouse cerebellum.** Report from published dataset^32^ against annotation from ABC mouse brain atlas.

Supplemental Data 2. **SAHA ann object of mouse cerebellum.** SAHA object from published dataset^32^ against annotation from ABC mouse brain atlas.

Supplemental Data 3. **HTML report expression of human PBMC.** Report from published dataset^33^ against annotation from COMBAT PBMC atlas.

Supplemental Data 4. **SAHA ann object of human PBMC.** SAHA object from published dataset^33^ against annotation from COMBAT PBMC atlas.

Supplemental Data 5. **HTML report expression of mouse isocortex.** Report from published dataset^35^ against annotation from ABC mouse brain atlas.

Supplemental Data 6. **SAHA ann object of mouse isocortex.** SAHA object from published dataset^35^ against annotation from ABC mouse brain atlas.

Supplemental Data 7. **HTML report expression of mouse spatial transcriptomics versus reference isocortex.** Report from published dataset^36^ against annotation from the isocortex of ABC mouse brain atlas.

Supplemental Data 8. **SAHA ann object of mouse spatial transcriptomics versus reference isocortex.** SAHA object from published dataset^36^ against annotation from the isocortex of ABC mouse brain atlas.

Supplemental Data 9. **HTML report expression of mouse spatial transcriptomics versus reference striatum.** Report from published dataset^36^ against annotation from the striatum of ABC mouse brain atlas.

Supplemental Data 10. **SAHA ann object of mouse spatial transcriptomics versus reference striatum.** SAHA object from published dataset^36^ against annotation from the striatum of ABC mouse brain atlas.

Supplemental Data 11. **HTML report expression of human microglial substates.** Report from published dataset^38^ against annotation from integrated glial atlas.

Supplemental Data 12. **SAHA ann object of human microglial substates.** SAHA object from published dataset^38^ against annotation from integrated glial atlas.

Supplemental Data 13. **HTML report expression of human astrocytic substates.** Report from published dataset^38^ against annotation from integrated glial atlas.

Supplemental Data 14. **SAHA ann object of human astrocytic substates.** SAHA object from published dataset^38^ against annotation from integrated glial atlas.

