## Supplementary figures and images for "Semi-automated annotation refinement accelerates cell type identification in brain spatial and single-cell studies"

### Supplemental Figures

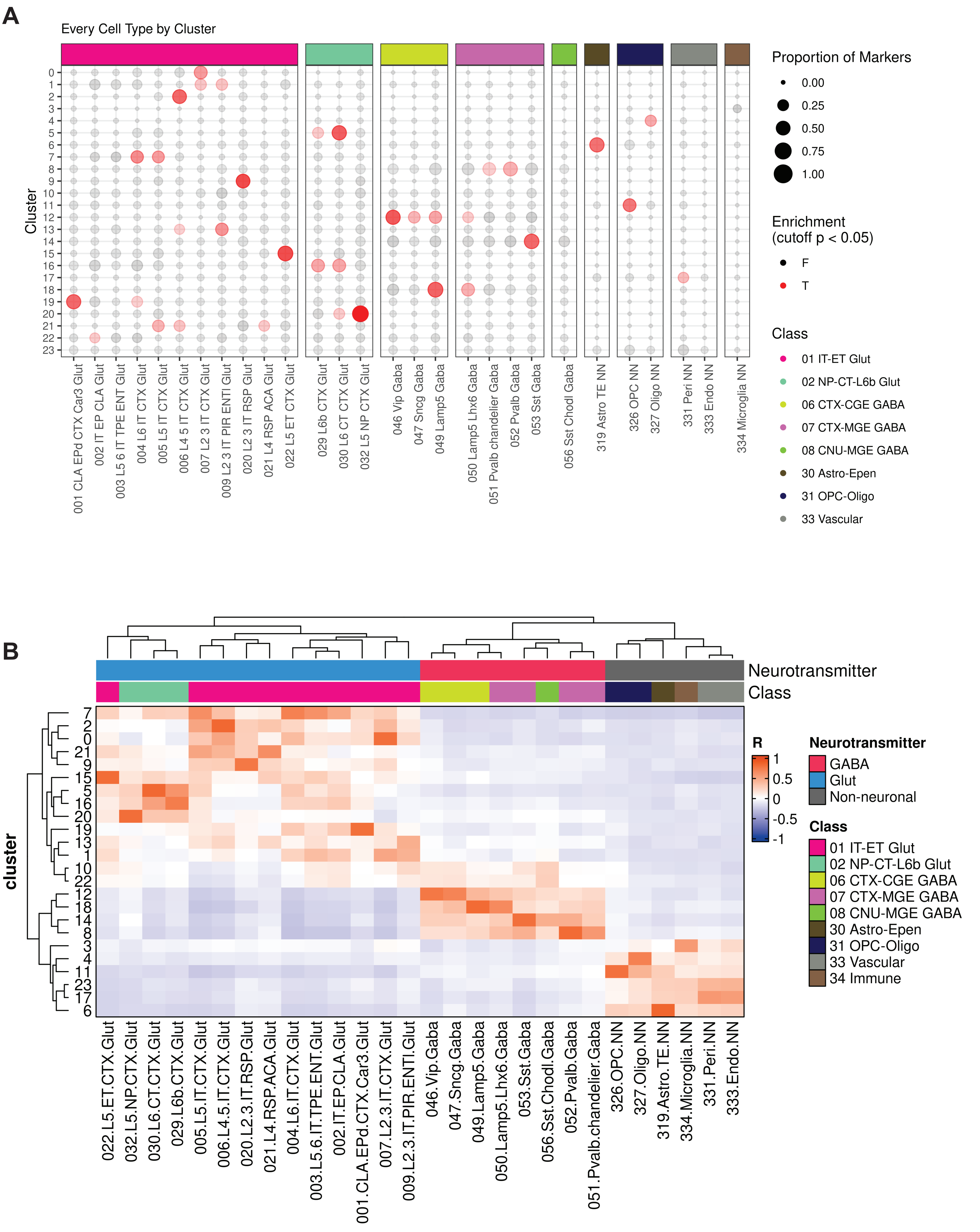

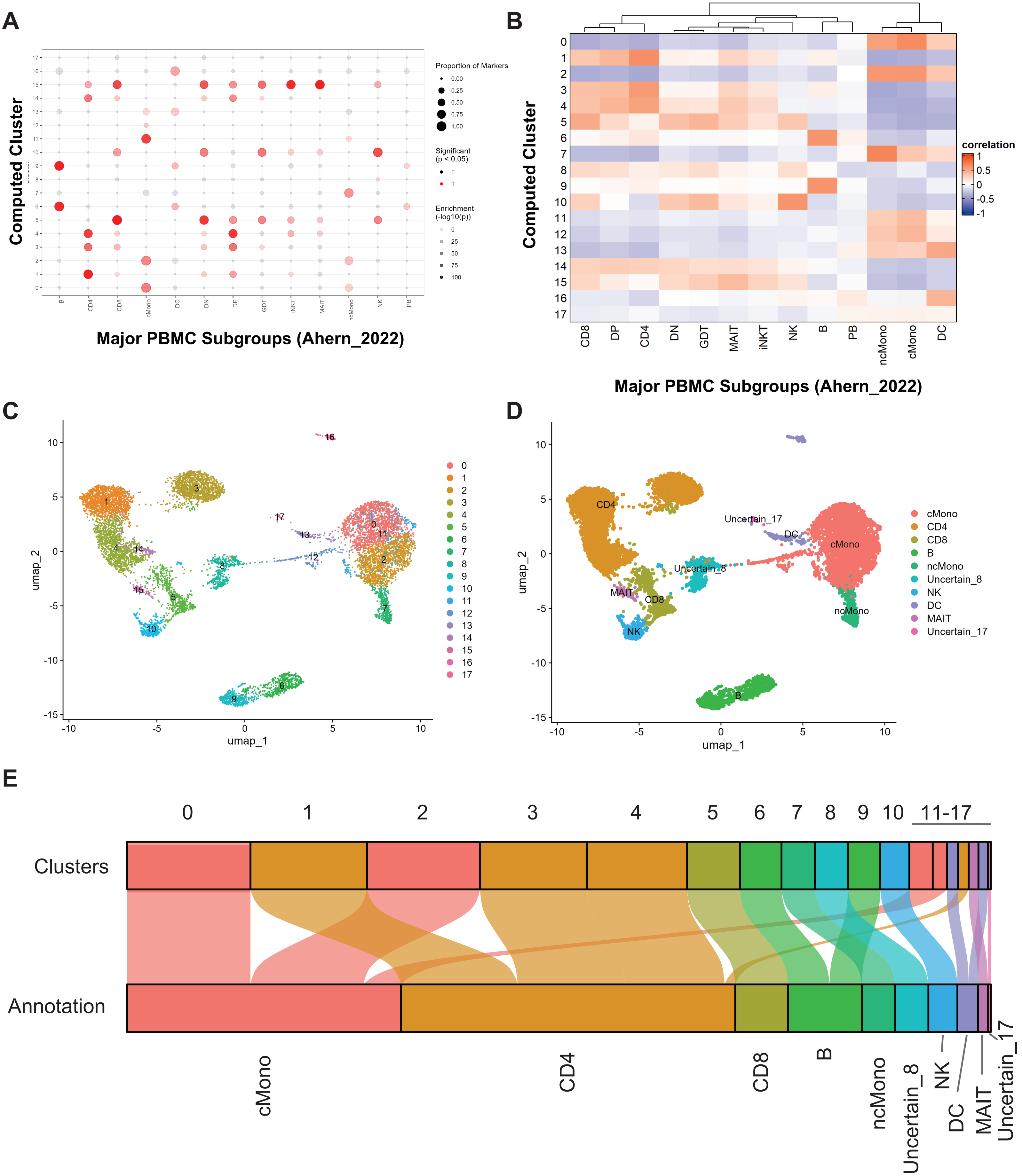

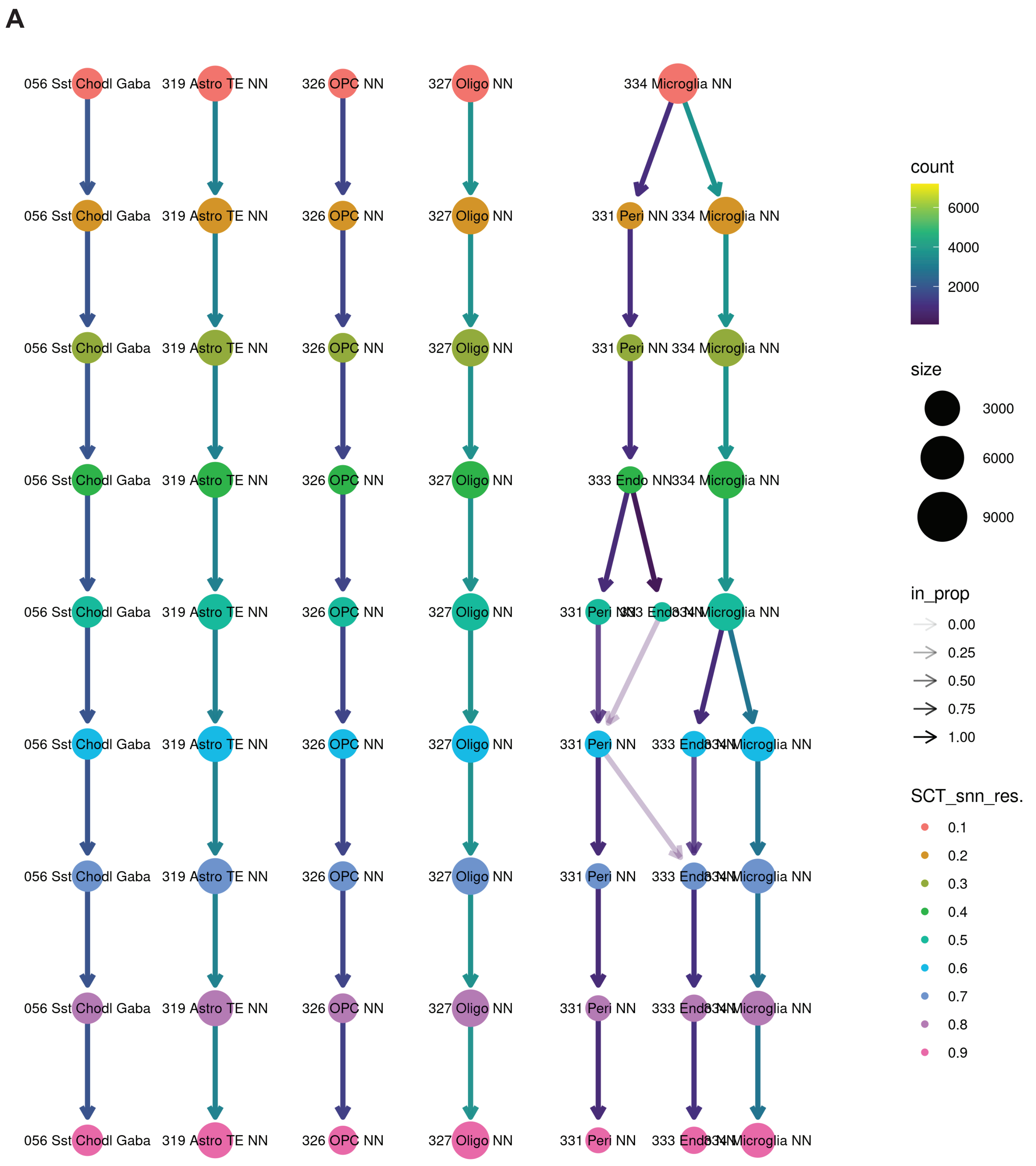

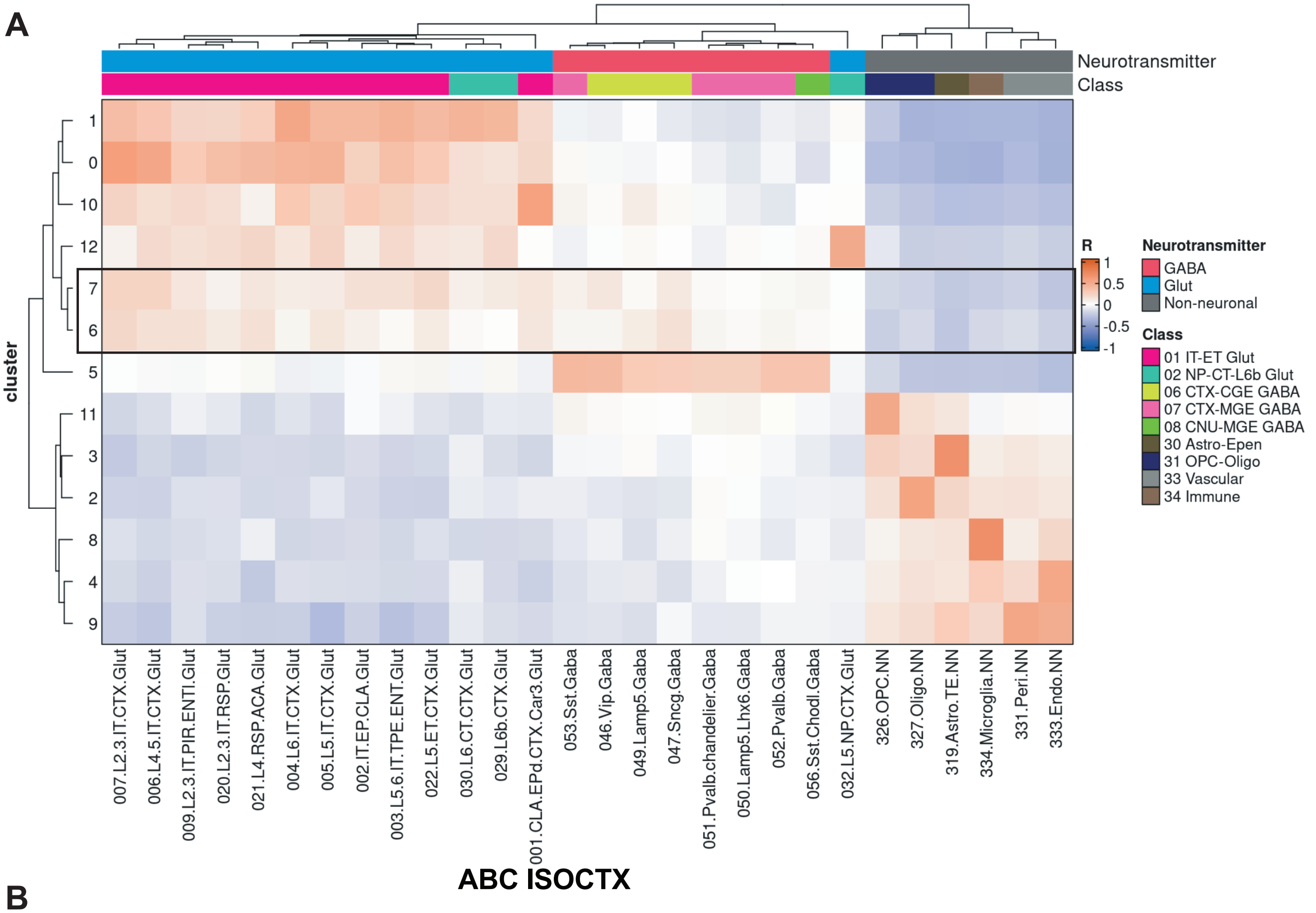

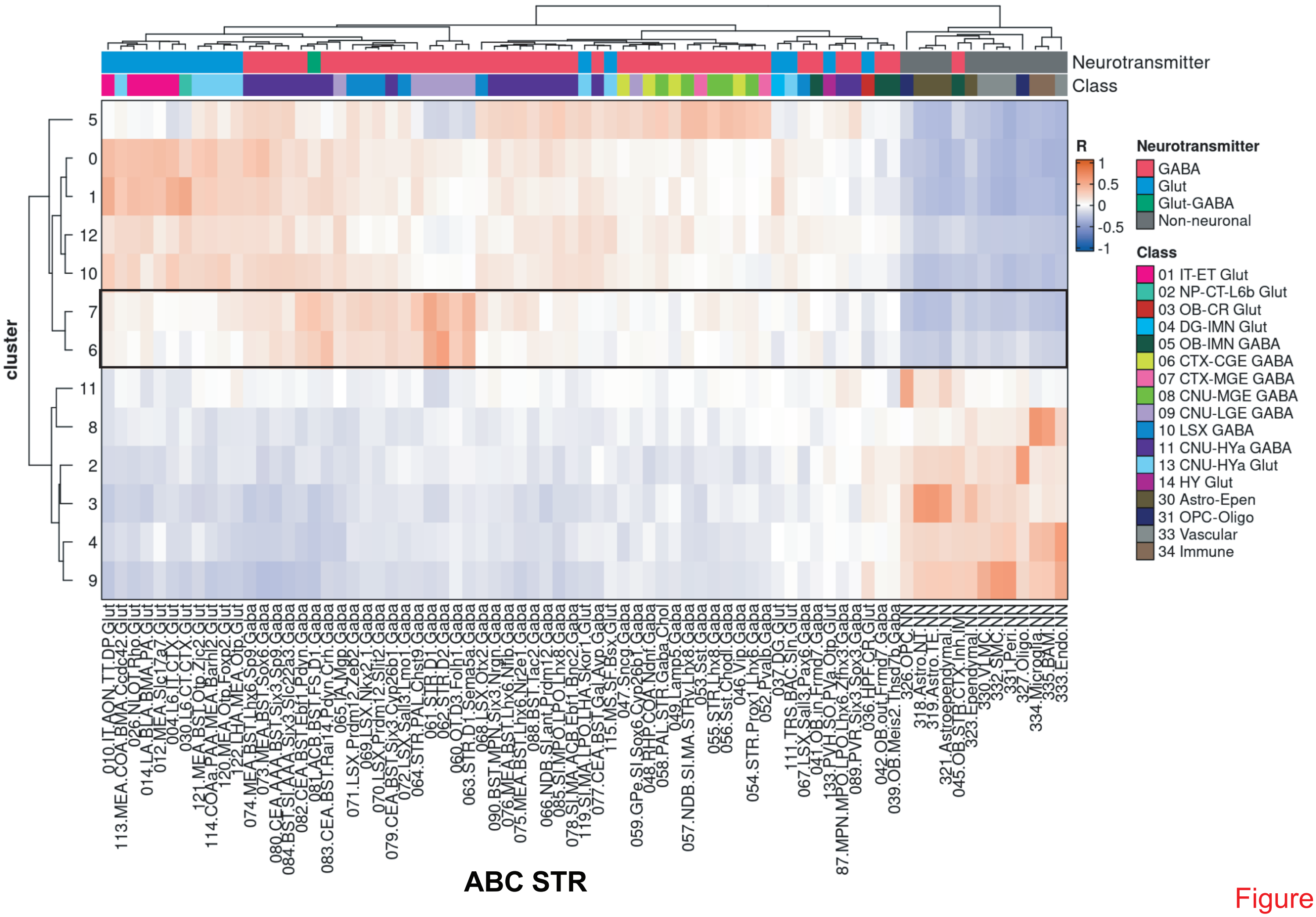

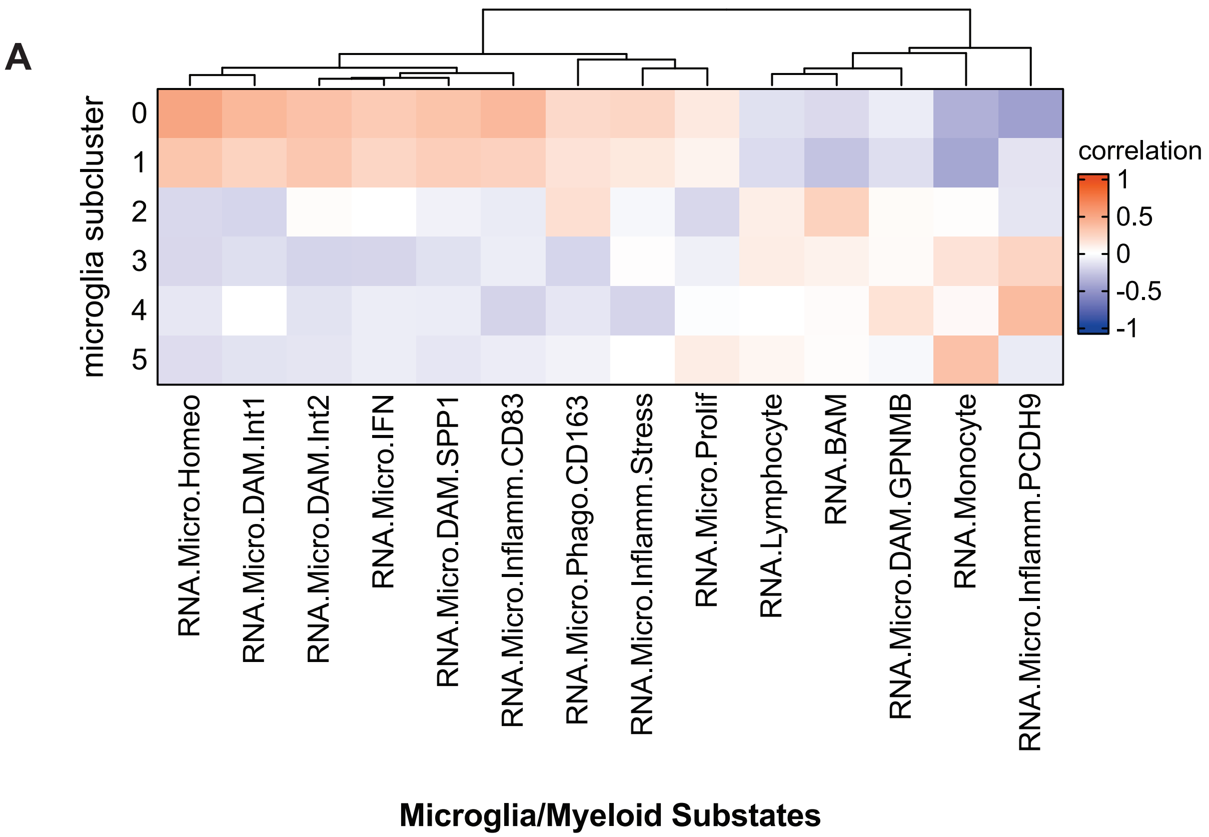

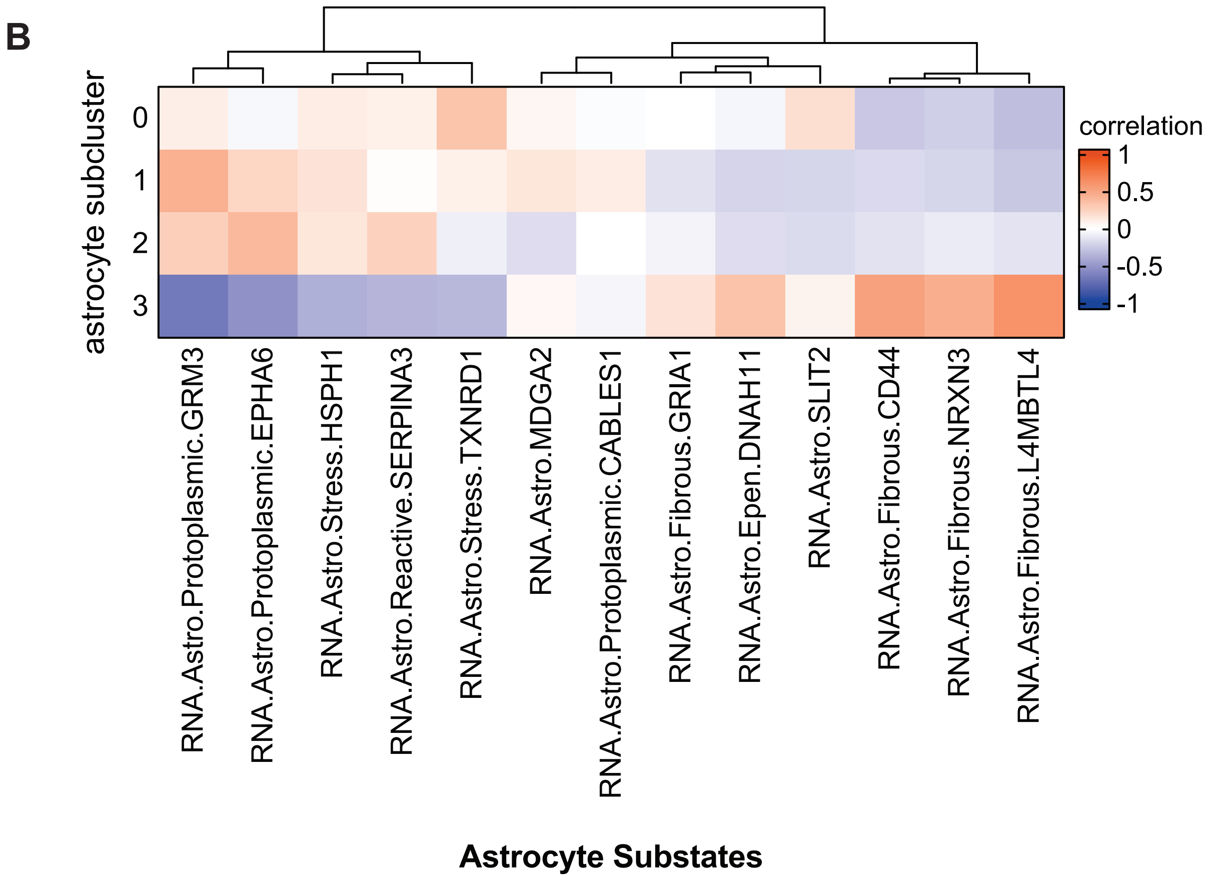
